# The Arabidopsis Concert of Metabolic Acclimation to High Light Stress

**DOI:** 10.1101/2023.02.14.528433

**Authors:** Gerd Ulrich Balcke, Khabat Vahabi, Jonas Giese, Iris Finkemeier, Alain Tissier

## Abstract

In plants, exposure to high light irradiation induces various stress responses, which entail complex metabolic rearrangements. To systematically study such dynamic changes, we conducted time course experiments from 2 minutes to 72 hours with *Arabidopsis thaliana* plants exposed to high and control light conditions. We performed comparative metabolomics, transcriptomics, redox proteomics and stable isotope labelling on leaf rosettes. Our data analysis identifies a set of synchronous and successive responses that provide a deeper insight into well-orchestrated mechanisms contributing to high light acclimation. We observe a downregulation of genes encoding light harvesting proteins and a transient restriction of genes involved in linear electron flow through photosystem I. C4 acids, produced via anaplerotic routes, strongly accumulate under high light conditions. Redox homeostasis is tightly balanced by reduced NADPH production, enhanced subcellular redistribution of reducing equivalents across several subcellular compartments via photorespiration and activation of processes that quench reactive oxygen species. In this well-orchestrated network, methylerythritol 2,4-cyclodiphosphate, fulfills a dual function as intermediate of plastidic isoprenoid production and as a stress signal molecule.

## Introduction

For plants, light is the primary source of energy for photosynthesis and a key factor in development. However, during their life span plants are exposed to fluctuating and variable light conditions. In particular, light intensity can dramatically change within a short time or for long periods, during which plants encounter light intensities that exceed their photosynthetic capacity (Mishra et al, 2012). In order to cope with those stressful situations, energy supply must be perfectly balanced with energy demand and redox metabolism must be fine-tuned at any time. Growth under low light conditions requires optimization of light absorption and energy transfer in order to use as much as possible of the available light energy. For example, under long term reduced light, algae and plants increase the size of their light harvesting pigment antennae for more efficient light capture (Muller et al, 2001). Furthermore, plants rely on photoreceptors such as phytochromes for shade avoidance responses and increase their stomatal aperture (Fiorucci & Fankhauser, 2017; O’Carrigan et al, 2014).

By contrast, under conditions that saturate the photosynthetic capacity, phototrophic organisms need to reduce absorption of light energy to avoid damage to their photosynthetic apparatus. Excess excitation energy ultimately leads to an increase in triplet chlorophyll, which is a potent photosensitizer of molecular oxygen forming singlet oxygen (Pospisil, 2016). High light irradiation exerts electron pressure on the electron transport chains of both photosystems, which in turn will affect the redox state in the chloroplast stroma, cytosol, and peroxisomes (Hashida et al, 2018; Lim et al, 2020; Steinbeck et al, 2020; Yokochi et al, 2021). When the activity of the photosynthetic electron transport exceeds the capacity of sink pathways that consume reducing equivalents, stroma over-reduction and sustained damage to the electron transport chains may result (Endo et al, 2005; Lima-Melo et al, 2019). Surplus reducing equivalents produced by linear electron flow can emerge in the form of strong reducing agents such as reduced thioredoxins or ferredoxins and NADPH with half-reduction potentials being low enough to transfer electrons to molecular oxygen (Asada, 2000; Kozuleva & Ivanov, 2016; Vogelsang & Dietz, 2020). Thus, high light stress rapidly leads to an increased formation of reactive oxygen species (ROS), which can cause oxidative damage to pigments, lipids, and proteins (Fichman et al, 2019).

In response to these stresses, plants have developed various acclimation processes, which together help to counter stromal over-reduction and ROS accumulation. High light stress prompts activation of superoxide dismutase, the gluthathione-ascorbate cycle, lipoxygenation and other direct or associated enzymatic reactions involved in ROS scavenging (Alsharafa et al, 2014; Asada, 2000; Xiao et al, 2012). Short-term protection against excess absorbed energy involves leaf avoidance movements and non-photochemical chlorophyll fluorescence quenching (NPQ), a process in which excess absorbed light energy is dissipated as heat (Foyer, 2018; Goss & Lepetit, 2015; Joliot & Johnson, 2011; Ruban, 2016). To rapidly regulate the accumulation of reducing equivalents and chloroplastic ATP production, phototrophic organisms control the fluxes through linear and cyclic electron flow (CEF) through the photosystems, with an increased share of CEF under high light (Walker et al, 2020). Under fluctuating light, CEF responds immediately to increased light intensity (Joliot & Johnson, 2011; Yamori et al, 2016). Excess electrons can be stored in starch or C4 carboxylic acids, be used for assimilation of sulfur or invested in the synthesis of strongly reduced carbon such as fatty acids or isoprenoids (Fernandez et al, 2017; Gibon et al, 2006; Kobayashi & DellaPenna, 2008; Kopriva et al, 1999; Medeiros et al, 2017).

In parallel, direct consumption of NADPH in the chloroplast is accomplished by light-dependent CO_2_ assimilation in the Calvin-Benson-Bassham cycle (CBB) (Michelet et al, 2013). Under ambient atmospheric CO_2_/O_2_ concentrations, the CBB enzyme ribulose-bisphosphate carboxylase (Rubisco) features, besides CO_2_ assimilation, high oxygenation activity and produces toxic 2-phosphoglycolate (2-PG). Recycling of 2-PG to 3-phosphoglycerate by photorespiration was long seen as a wasteful process because formally it reduces the CO_2_ assimilation rate and consumes ATP (Fernie & Bauwe, 2020). However, intrinsically tied to the dual activity of Rubisco, this recycling pathway forms a link between central carbon and nitrogen metabolism and shuttles excess reducing power across four subcellular compartments (Nunes-Nesi et al, 2010).

The plastidial methylerythritol phosphate (MEP) pathway provides precursors of several isoprenoids that play essential roles in photosynthesis and protection against high light such as chlorophylls, carotenoids, plastoquinones or tocopherols (Brunetti et al, 2015). As a result, its regulation may be crucial in the adaptation to high-light stress. In addition, the MEP pathway represents a potential sink to take up excess reducing power because three of its seven reactions are reductions that consume NADPH and reduced ferredoxins (Figure-SI 1). Furthermore, accumulation of the small molecule intermediate 2-C-methyl-D-erythritol-2,4-cyclodiphosphate (MEcPP) plays an important part in retrograde signaling of perturbed redox conditions towards the activation of nucleus-encoded genes involved in the defense against ROS (Jiang et al, 2020; Xiao et al, 2012). Therefore, regulation of the MEP pathway under high light conditions could be of crucial importance for the acclimation process. Sensing and mediation of oxidative stress under high light is also supported by assembling chloroplasts and nuclei in close vicinity, where MEcPP, other retrograde signaling molecules, and H_2_O_2_ itself may induce rapid transcriptional stress responses directly (D’Alessandro et al, 2018; Exposito-Rodriguez et al, 2017; Foyer, 2018; Ramel et al, 2012).

Thus, the response to high light irradiation in plants entails complex metabolic rearrangements, which were studied previously in some detail. As of late, newly emerging ‘omics techniques allow broader analysis of such metabolic adaptations and aim to connect detailed results with a systems perspective on plant acclimation to high light (Kudo et al, 2019; Lee et al, 2020; Moreno et al, 2021; Schmitz et al, 2014; Treves et al, 2020).

Comprehensive analyses of the dynamic interplay between individual stress responses to high light were performed recently, using either RNA-seq data only (Alvarez-Fernandez et al, 2021; Huang et al, 2019), proteomics only or by combining several ‘omics techniques. Yet, those focused on ectopic expression mutants (lycopene *β*-cyclase1) or specifically on the accumulation of phenylpropanoids (Moreno et al, 2021; Neugart et al, 2016; Niedermaier et al, 2020). However, in depth studies of dynamic responses to light stress that link transcriptional responses to changes directly seen at the metabolite level in central carbon and secondary metabolism are so far lacking.

Here we combine transcriptomics, deep metabolomics, redox proteomics, and isotope labeling to study coordinated responses in *Arabidopsis thaliana* stressed with high light. In order to analyze dynamic responses, leaf rosettes were sampled over the course of minutes to three days. For a broad metabolite profiling of central carbon, energy and plant secondary metabolism we use a set of targeted and untargeted methods and apply molecular clustering techniques (Treutler et al, 2016).

## Results

### Experimental Design and Phenotypic Observations

Rosettes of *Arabidopsis thaliana* ecotype Columbia (Col) were exposed to two different light conditions and were harvested at eight time points ranging from 2 min to 72 h (Figure 1A-C). We consider the time points until 300 min as the early response phase, while the time points from 24 to 72 h, which were interrupted by nights represent the acclimation phase. Dynamic metabolic, isotopic, redox-proteomic, and transcriptional responses to high light stress were profiled by LC-MS/MS and RNA-seq, respectively. Altogether, 186 metabolic features targeting the central carbon and energy metabolism (CCEM), 23,199 features of untargeted metabolomics, the expression of 23,658 genes and the thiol-group oxidation state of 140 selected proteins formed the data basis for further analysis (Data-SI 1, Data-SI 2, Table-SI 1). During this period, plants under high light showed no apparent symptoms of photo damage, but gradually accumulated anthocyanins (Figure 1 D-E) indicative of a stress response to high light.

**Figure 1.**
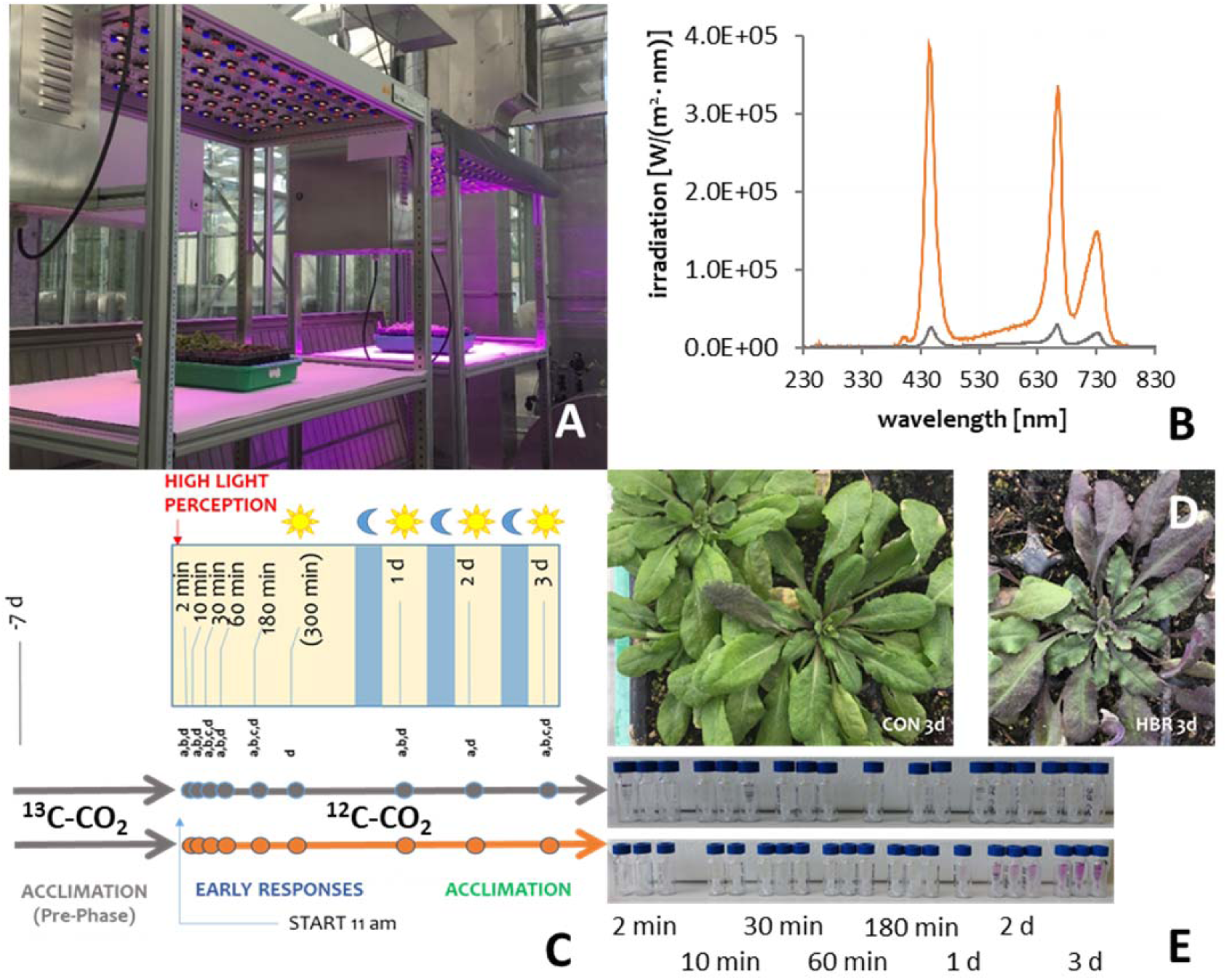
Experimental Setup and phenotypic observations during high light stress in Arabidopsis thaliana. **A.** Experimental setup using Rhenac LED panels, with 130 and 1300 µmol m^−2^ s^−1^, respectively. **B**. High and control light irradiance spectra used in these experiments. **C**, Sampling design and day/night conditions for two experiments: one without^a,b,c^ and one with pulse-chase isotopic labeling^d^, (sampling points for ^a^metabolomics, ^b^transcriptomics, ^c^redox-proteomics, ^d^isotopic labeling). **D**, Phenotypic observations after 3d of control light (CON) and high light (HBR). E, Progressive accumulation of anthocyanins in aqueous extracts in the course of control and high light irradiation. Grey and orange lines indicate control light and high light conditions, respectively.

Isotopic enrichment in small molecules with stable carbon isotopes allows to estimate metabolic fluxes in response to high light (Ma et al, 2014). Here, in a second experiment, we opted for reverse ^13^C labeling because it allows light stress experiments to be performed under ambient natural CO_2_ levels, which in turn permits rapid arrest of metabolism in plant leaves when harvesting. After a 7-day long ^13^C labeling period preceding the actual light stress experiment plants were then exposed to either control or high light and simultaneously to ambient carbon dioxide levels of 450-550 ppm ^12^C-CO_2_ (chase phase, Figure 1C). High enrichments with ^13^C carbon were achieved for all metabolites of the central carbon metabolism, which is demonstrated by low ^12^C fractional isotope enrichment (FIE) at the starting point of the chase phase (Figure 5).

### Dynamic Responses of Transcriptome, Metabolome, Carbon Flux, and Redox-Proteome in the Course of High Light Irradiation

Pairwise comparison between light-stressed and control light samples identified differentially expressed genes (DEGs) and differentially abundant metabolites (DABs) at each time point (Table-SI 2, Data-SI 1). Further analysis indicated substantial differences in the time course of DEGs and DABs. As a response to light stress, the number of DEGs at each time point increased mostly from 0-180 min (Figure 2A). After 24 h, 24% of all detectable transcripts showed differential regulation between high and control light. Interestingly, after long-term exposure to high light the number of DEGs showing downregulation deceased whereas the number of upregulated genes continually increased until 72 h (Figure 2A). By contrast, levels of many metabolites showed increased abundances not before 24 h indicating overall an enhanced biosynthetic activity within the scale of several hours to days after exposure to high light. Pairwise clustering of transcript and metabolite profiles reflected this as transcripts segregated into two distinct clusters for early time points (2 and 10 min and 30 to 180 min) whereas a large number of metabolites showed pattern changes much later (Figure 2B). K-means time clustering of transcriptional regulation disclosed 15 clusters of which four selected clusters are presented in Figure 2 panel C (all 15 in *Table-SI 3*).

**Figure 2.**
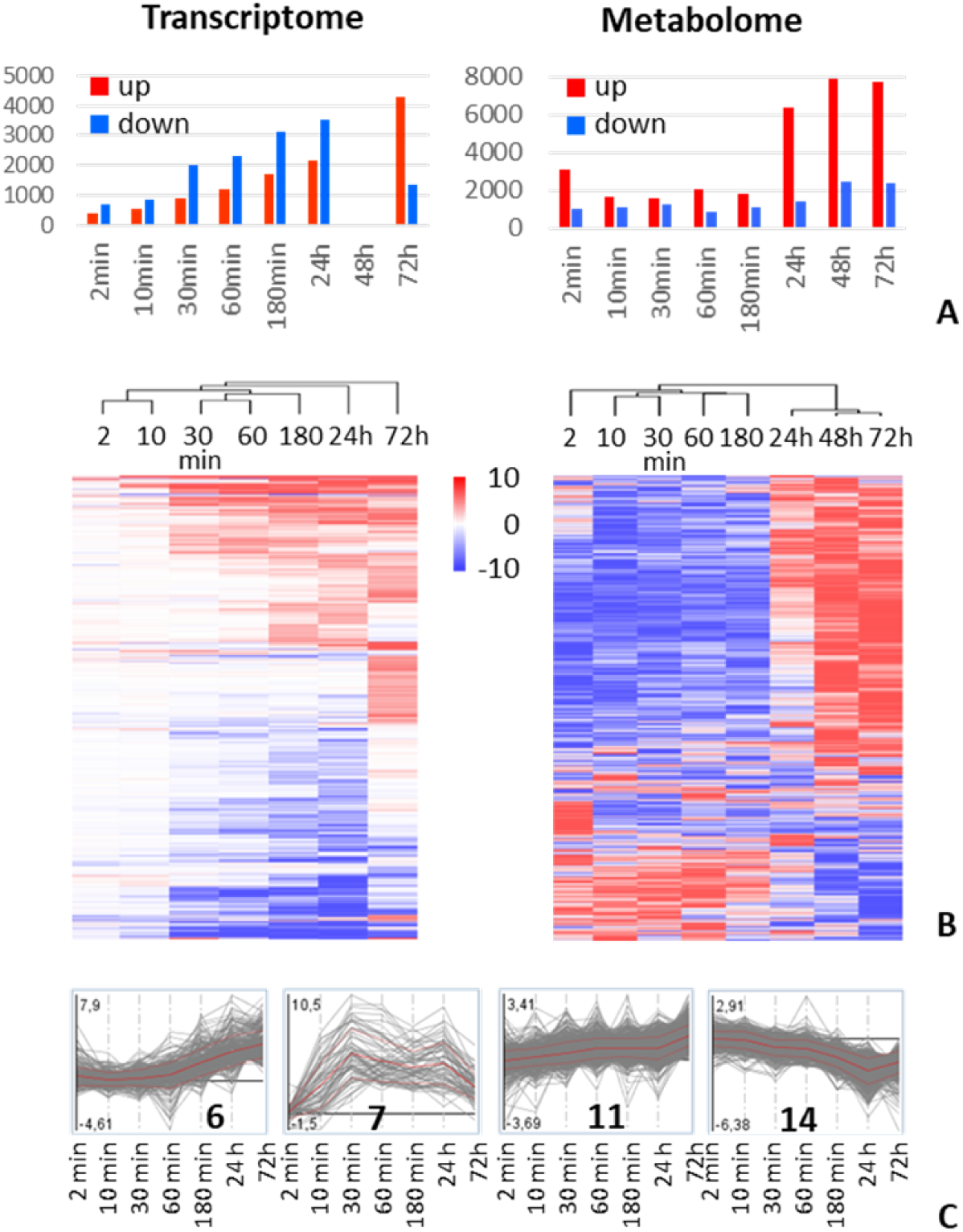
Analysis of differentially expressed genes (left) and differentially abundant metabolites (right). **A,** Total number of differentially regulated signals at each time point, filters: −1 ≥ log_2_FC ≥ 1, p ≤ 0.05 signal threshold ≥ 0.5 TPM for transcripts, ≥10% or average intensity for metabolites. **B,** Time cluster analysis, filters: −1 ≥ log_2_FC ≥ 1, p ≤ 0.05, signal threshold ≥ 5 TPM for transcripts, ≥10% or average intensity for metabolites. **C,** K-means cluster analysis of co-regulated genes showing 4 out of 15 time clusters of the transcriptomics data.

#### Very early responses to high light irradiation

Differential gene expression within the first 10 min of light stress was observed only for a small number of genes. Among those were transcripts encoding Ferredoxin 1 *(FD1,* AT1G10960), early light-inducible protein (e.g. *ELIP1*, AT3G22840) but also mitochondrial ubiquinol-cytochrome C reductase (AT5G25450) (Figure 3 panels A,C,M, Table-SI 4). While early regulations of *FD1* and *ELIP1* do not come as a surprise – both plastid enzymes have functions in the maintenance of redox homeostasis and avoidance of photo-damage – rapid activation of mitochondrial respiration raises the question of the origin of the carbon source for increased oxidative phosphorylation (Hutin et al, 2003; Kozuleva & Ivanov, 2016).

**Figure 3.**
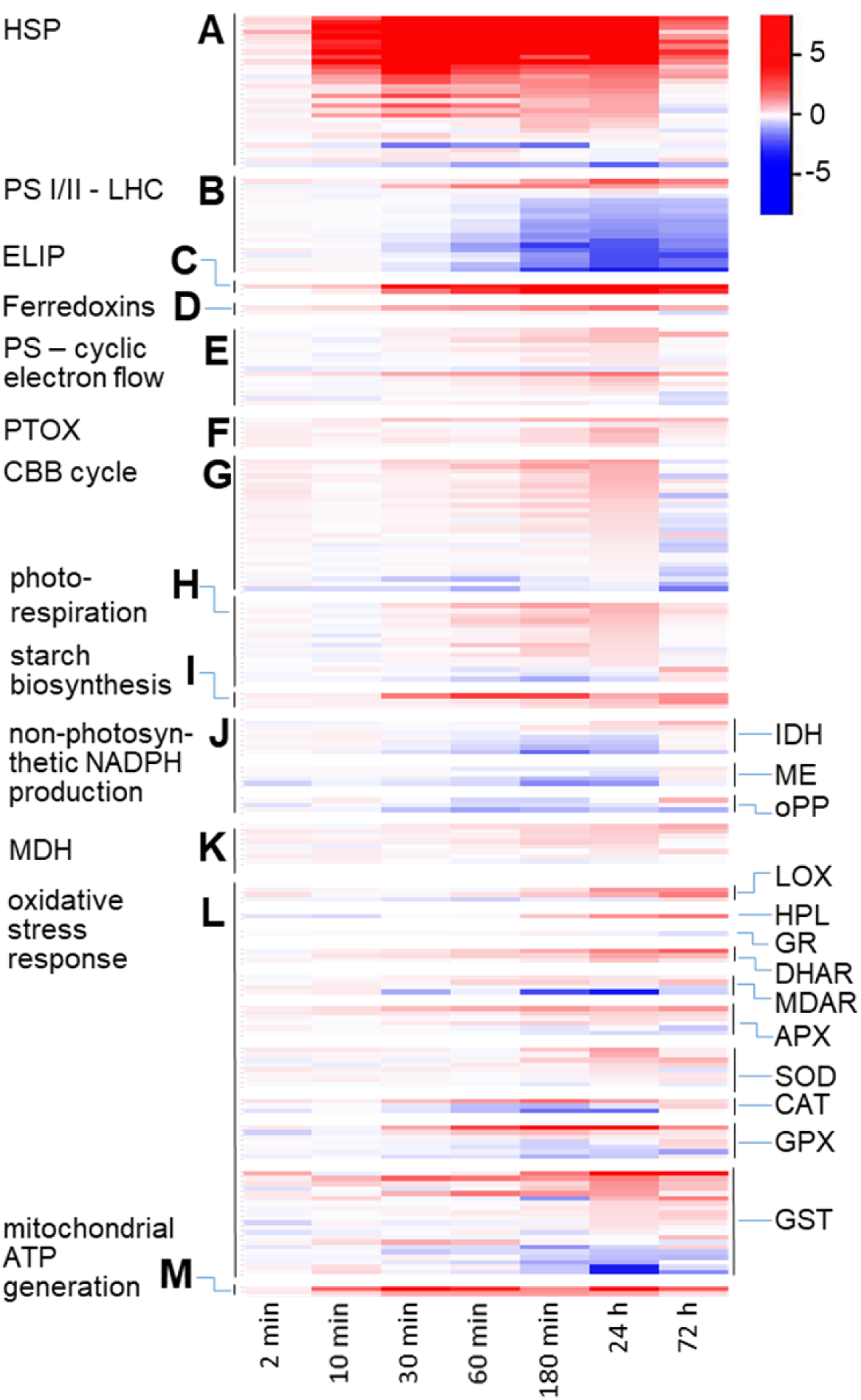
Differential expression profile within selected gene groups. **A**, Heat shock proteins. **B**, Photosystem I and II – Light harvesting complexes. **C**, Early light inducible protein, **D**, Ferredoxins. **E**, Photosystems – Cyclic electron flow. **F**, Plastid terminal oxidase. **G**, Calvin-Benson-Bassham cycle. **H**, Photorespiration. **I**, Starch biosynthesis. **J**, Non-photosynthetic NADPH production. **K**, Malate dehydrogenases. **L**, Oxidative stress response. **M**, Mitochondrial ATP production. The graph depicts log_2_-fold changes between high light and control light (n=3) at a given sampling time (all data are unscaled).

GO enrichment analyses of most early responding genes (upregulation after 2 min of light stress) shows over-representation of genes involved in response to hydrogen peroxide, to reactive oxygen species and heat (Figure 2C Cluster 7, Figure-SI 2). Accordingly, heat shock proteins (HSP) showed strong upregulation under light stress within the scale of minutes. HSP reached the highest differential expression versus normal light controls between 30 min and 24 h and declined again towards 72 h of light stress (Figure 3A). Induction of HSP upon light stress has been observed by others and was interpreted as an overlap between heat stress and high light responses (Huang et al, 2019). Therefore, we investigated the heat contribution of the LED light source. We measured the temperature in the leaf layer and detected only a moderate increase by 2.5 °C in light-stress panels versus control panels indicating that LED heat emission did not strongly contribute to the severe upregulation of HSP.

A rapid increase in metabolite levels and faster isotopic labeling in intermediates of the CBB cycle in light-stressed plants versus plants grown in normal light conditions was observed already after 2 min (Figure 4A, Figure 5). Likewise, metabolic levels of the glycolytic, photorespiratory and MEP pathway intermediates instantly increased upon high light exposure (Figure 4B,C,E). However, faster ^12^C-FIE in sugar phosphates as well as in 6-phosphogluconate, phosphoenolpyruvate, photorespiratory and MEP pathway intermediates was observed after high light exposure. The strongest increase in ^12^C-incorporation was observed for the N+0 isotopolog, which only represents fresh carbon in metabolites being *de novo* synthesized (Figure-SI 3).

**Figure 4.**
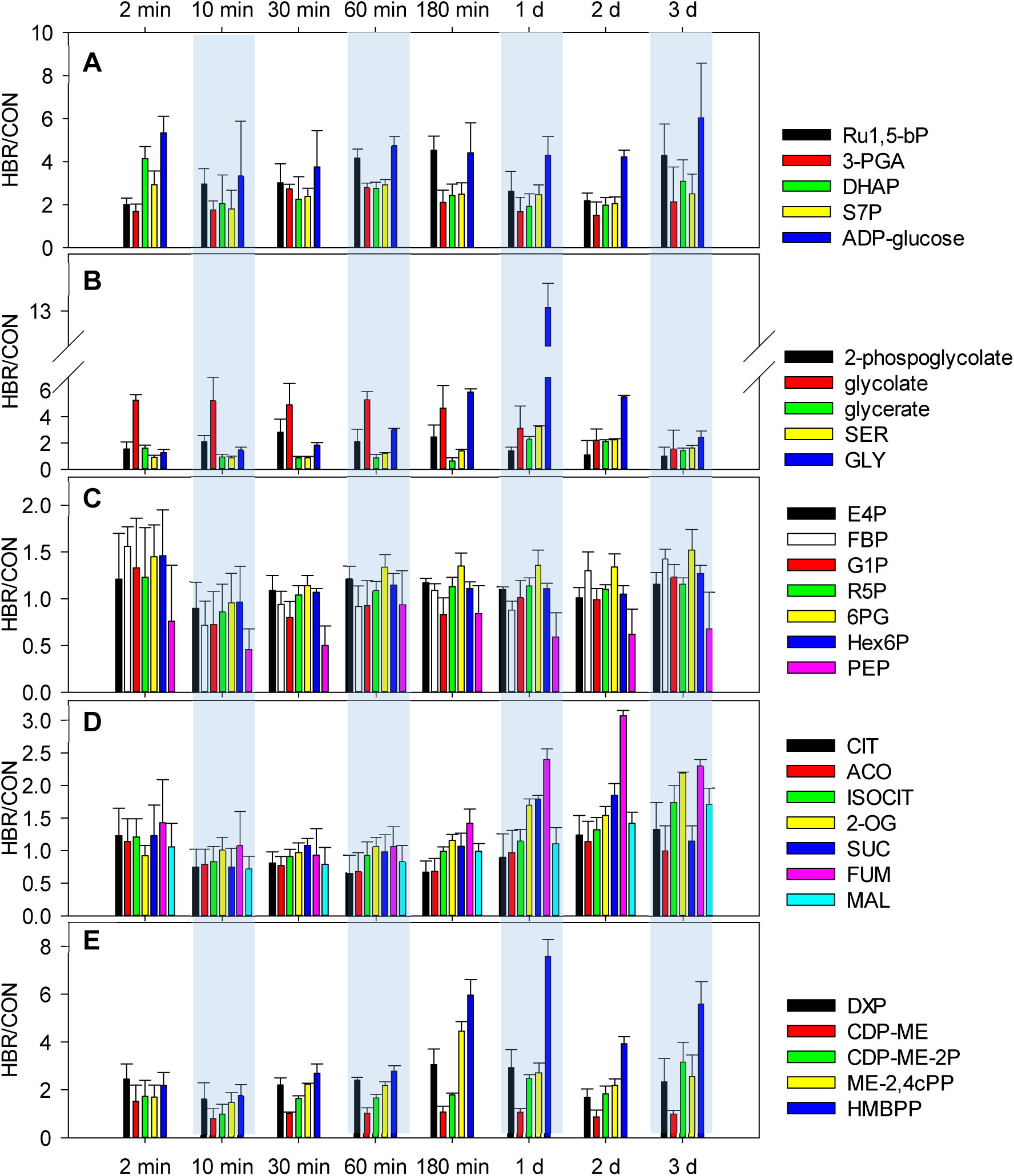
Central carbon metabolism: relative metabolite signal intensity after exposure to high light (HBR) versus normal light (CON). **A**, Calvin-Benson-Bassham cycle and starch biosynthesis. **B**, Photorespiration. **C**, Sugar metabolism. **D**, TCA cycle. **E**, Methylerythritol phosphate pathway. Structural identifiers for metabolite abbreviations are given in **Table-SI 5**.

**Figure 5.**
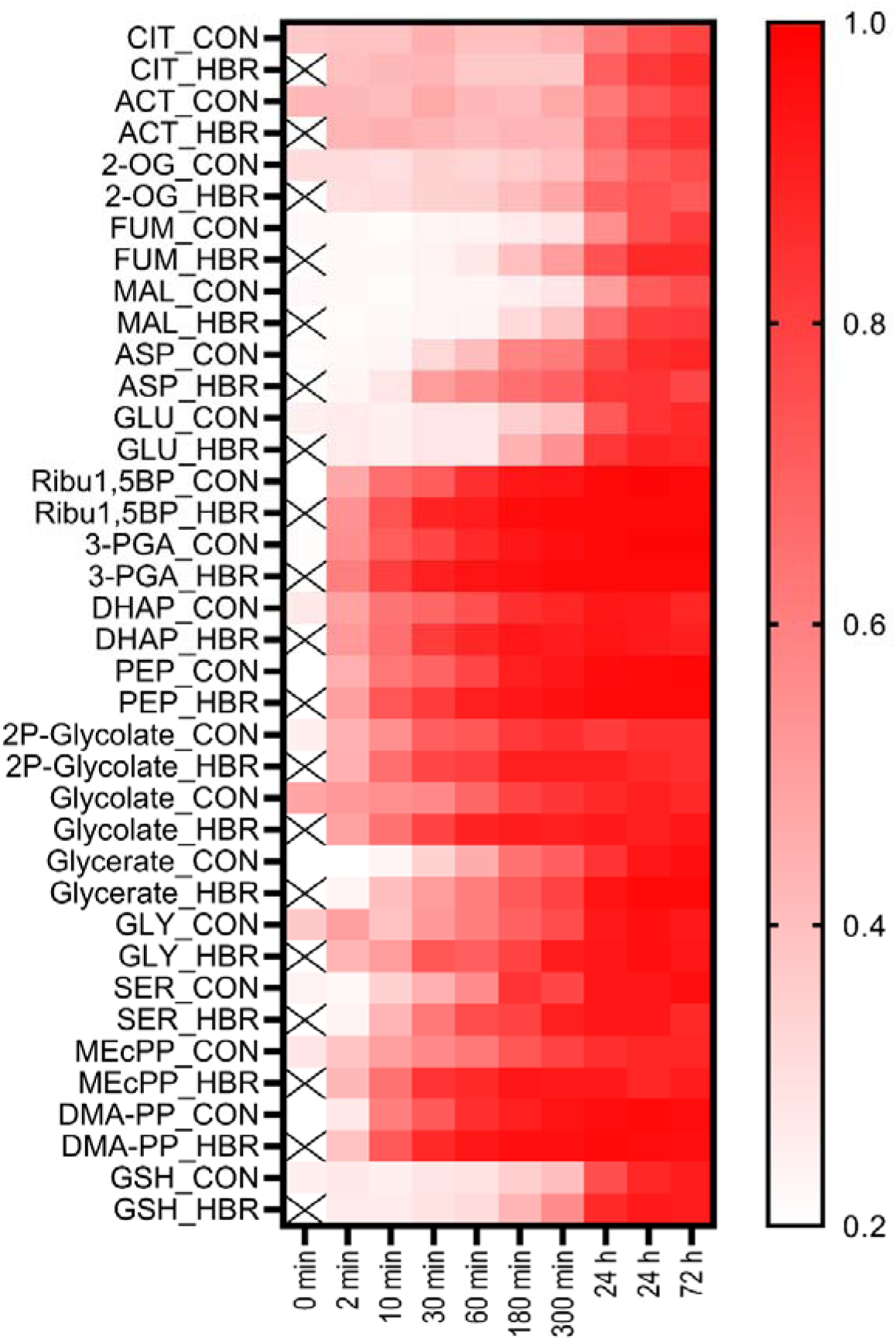
Fractional isotopic enrichment with ^12^C after pulse-chase labeling with CO_2_ under control (CON) and high light (HBR). Structural identifiers for metabolite abbreviations are given in Table-SI 5.

#### Upon light stress photosystems, dark reactions and central sugar metabolism exhibit different regulatory patterns

In contrast to HSP, transcripts encoding light harvesting complexes of both photosystems were slowly repressed in light-stressed plants relative to the control (Figure 2C cluster 14, Figure 3, gene group B). Simultaneously, the expression of genes encoding components of the cyclic electron flow (CEF) and plastoquinone terminal oxidases (PTOX) increased from 180 min to 24 h (Figure 3, gene groups E-F). Concomitantly, transcripts encoding enzymes of the CBB cycle, photorespiration and starch biosynthesis showed significantly increased expression under light stress from 30-180 min to 24 h (Figure 3, gene groups G-I). Notably, after 72 h of light stress most transcripts within the categories “PS-cyclic electron flow, PTOX, CBB cycle and photorespiration” decreased to values only slightly higher than control light samples or were indistinguishable from them, whereas transcripts for starch biosynthesis remained elevated under high light (Figure 3, gene groups G-I, Table-SI 4).

Metabolites of the CBB cycle increased already within 2 min of light stress and remained at higher levels in light-stressed samples relative to their controls throughout the experiment (Figure 4A). A similar trend was observed for ADP-glucose, which is an intermediate of starch biosynthesis (Figure 2C cluster 11, Figure 4A). Thus, the metabolite levels in these pathways increase before the transcript levels and remain high despite transcript levels returning to lower levels (Figure 3, gene groups G,I). This indicates a coordinated but independent regulation of metabolite and transcripts.

Similarly, several intermediates involved in photorespiration, such as 2-phosphoglycolate, glycolate or glycine, showed rapidly increased levels within minutes after perception of high light (Figure 4B). However, 2-phosphoglycolate and glycolate levels then decreased to levels comparable to normal light conditions (day 1 to day 3), whereas their downstream intermediates glycerate, glycine and serine accumulated in light stressed plants with a maximum at day 1 and then returned to moderately increased levels.

Over the course of the experiment sugar phosphates and 6-phosphogluconate (6PG), which is an intermediate of the oxidative pentose phosphate pathway, showed constantly increased values in high light versus normal light (Figure 4C). Among the glycolytic intermediates, phosphoenolpyruvate (PEP) alone showed a distinct behavior with slightly decreased but constant levels under high light throughout the experiment (Figure 4C). This indicates a specific role of PEP in response to high light, compared to other glycolytic intermediates. However, isotopic labeling revealed faster ^12^C-incorporation into PEP in high light than in low light indicating increased flux through PEP during high light (Figure 5, Figure-SI 3).

In agreement with the nearly constant levels of glycolytic intermediates, most transcripts encoding glycolysis, phosphoenolpyruvate carboxylase and carboxykinase enzymes showed only little change during acclimation to high light (Table-SI 4). Taken together, light stress induces the downregulation of genes involved in light harvesting while upregulating genes involved in CBB cycle carbon fixation, photorespiration, and starch biosynthesis during the early phase. Stable levels of many sugar phosphates and their biosynthetic genes reflect homeostasis despite different light conditions.

#### Light stress leads to reduced expression of NADPH producing processes

Simultaneous downregulation of the light harvesting complex (LHC) and upregulation of the CEF and PTOX in the photosystems are seen as mechanisms to counteract over-reduction in the chloroplast stroma induced by accumulation of strong reductants such as reduced ferredoxins and NADPH. In addition to photosystem I, isocitrate dehydrogenase (IDH), malic enzyme (ME), and the oxidative branch of the pentose phosphate pathway (OPP) can generate NADPH. Under light stress, all of these NADPH producing routes are transcriptionally repressed starting from 30 min onwards (Figure 3, gene group J). After 72 h of light stress, only cytosolic IDH (AT1G65930) and one isoform of glucose 6-phosphate dehydrogenase (AT5G40760) remained repressed whereas most other genes in group J encoding ME, IDH and OPP showed similar or even elevated gene expression under high light (Table-SI 4). Thus, coinciding with the transcriptional downregulation of photosynthetic reactions producing energy-rich reducing equivalents, alternative routes producing NADPH are also downregulated within the first day of high light exposure.

#### Enhanced accumulation of C4 acids under high light is driven by anaplerotic routes

Metabolite profiling of TCA cycle intermediates showed increasing accumulation of malate (MAL) and fumarate (FUM) within the scale of several hours to days (Figure 4D). By contrast, citrate and aconitate did not show altered levels at all (Figure 4D). Reverse labeling after a ^13^C-enrichment phase show much slower incorporation of fresh, i.e. ^12^C, carbon into TCA cycle intermediates than observed for sugar phosphates (Figure 5). Individual acids display very different labeling kinetics, which are not only affected by light intensity but also showed differences after a night period (day 1-3). Despite their high abundance, the pool of the N+0 isotopolog for FUM and MAL increased much more rapidly and more intensively under light stress conditions as compared to control light. By contrast, within 300 min the FIE and patterns of the isotopic distribution of CIT and aconitate (ACO) do not differ from the pattern observed at 0 min, at the start of the chase phase (Figure 5, Figure-SI 3). Unlike citrate and aconitate, 2-oxoglutarate (2-OG) displays a very small pool size in *Arabidopsis* leaves (Szecowka et al, 2013). Therefore, clockwise carbon flow across the TCA cycle should result in rapid labelling of 2-OG. However, chase labeling showed only little synthesis of the N+0 isotopolog in 2-OG from 0-300 min compared to the levels observed after the first night and noteworthy incorporation of ^12^C was observed only after several hours. Similar observations were made for glutamate, which shares the same carbon body with 2-OG. Together, these strongly differential labelling patterns between C4 acids on the one hand and C5/C6 acids on the other indicate that during the day there is little clockwise activity in the TCA cycle from CIT to 2-OG. By contrast, the formation of C4 acids is strong during daylight and even increases upon exposure to high light. Thus, as C4 is not made from citrate, anaplerotic formation of MAL and FUM via PEPC is anticipated, which is seen as a way to balance redox homeostasis under high light (Igamberdiev & Eprintsev, 2016; Tcherkez et al, 2009). Enhanced ^12^C-enrichment in FUM, MAL, and in ASP and PEP under light stress as well as strong transcriptional upregulation of chloroplastic glucose 6-phosphate translocator 2 (GPT2, up to 42 fold) infer enhanced export of sugars phosphates, cytosolic glycolysis and formation of oxaloacetate via high activity of phosphoenolpyuvate carboxylase (PEPC) (Figure 5,Table-SI 4). We show here that this flux plays a key role in the defense against high light.

In Arabidopsis, citrate (CIT), MAL and FUM - due to high vacuolar accumulation – can reach micromoles per gram fresh weight (Szecowka et al, 2013). To estimate the contribution of the most abundant compounds of the TCA cycle to carbon storage, we performed quantitative assays for FUM, MAL, and CIT and compared their values per C1 carbon units to the accumulation of starch (Figure 6). Under high light, starch levels increase, supported by transcriptional upregulation of biosynthetic genes (Figure 2C, cluster 11, Figure-SI 2). However, direct comparison shows that the amount of reduced carbon that is stored in FUM and MAL is by far not negligible. In control light conditions, the cumulated amount in moles of C of FUM, MAL, and CIT was 11.2 fold that of starch, and under high light stress it was still 3.4 fold higher. Dramatic increases were noted for FUM (2.1 fold), MAL (2.9 fold), and starch (6.8 fold), while CIT showed only a non-significant marginal increase when comparing high and low light conditions. Thus, photosynthetic carbon storage in form of MAL and FUM in vacuoles represents a so far underestimated way of carbon storage besides the formation of starch in chloroplasts.

**Figure 6.**
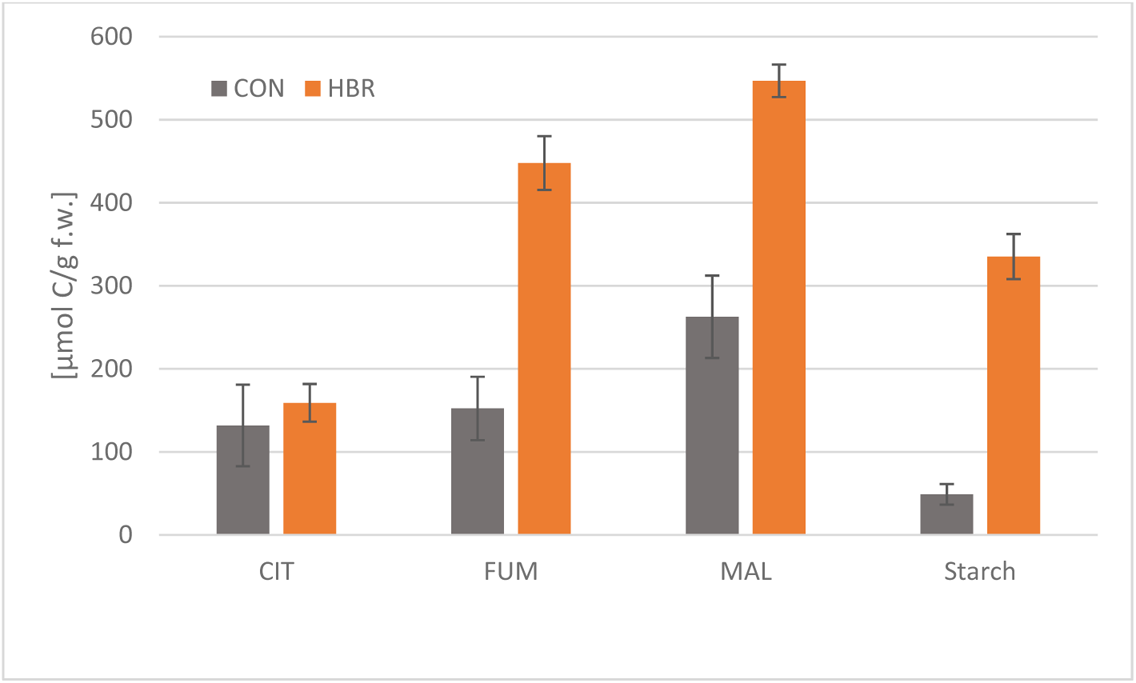
Absolute quantification of malate, fumarate, citrate, and starch. Samples were collected at 11 am after 3 d of treatment (n=4 plants). CON: control light; HBR: high light.

#### Redox homeostasis is maintained under high and normal light

Next, we sought to understand how light stress influences the redox state in the rosette material. We observed that genes involved in ROS-counteracting processes such as the glutathione-ascorbate cycle, superoxide dismutases, lipoxygenation, hydroperoxide lyase and glutathione peroxidases as well as transferases are transcriptionally upregulated after 10 min of high light and onwards (Figure 3, gene group L, Figure-SI 4, Table-SI 4). For example, we found transcriptional upregulation of cytosolic (AT1G07890, up to 4-fold) and stroma-localized ascorbate peroxidase (APXS, AT4G08390 up to 2-fold). Upregulation remained high until the last sampling point at 72 h.

Redox-sensitive metabolites were assayed in their oxidized and reduced forms to check for their levels and ratios during light stress (Figure 7). Couples of oxidized and reduced glutathione (GSSG/GSH) and NAD(P)+/NAD(P)H revealed maintenance of redox homeostasis over the first 30 min after induction of light stress followed by a gradual accumulation of only the oxidized forms within the scale of hours to days. The accumulation of GSSG under high light was followed by an increasing accumulation of ascorbate (ASC) from 1 to 3 days. Notably, no strong increase in NADPH or GSH was observed throughout the experiment. NAD^+^, NADP^+^ and GSSG are known to be very weak oxidants and ASC is only a mild reductant. Therefore, accumulation of these redox-sensitive metabolites is seen as a consequence of detoxification reactions such as the glutathione-ascorbate cycle (Figure-SI 4).

**Figure 7.**
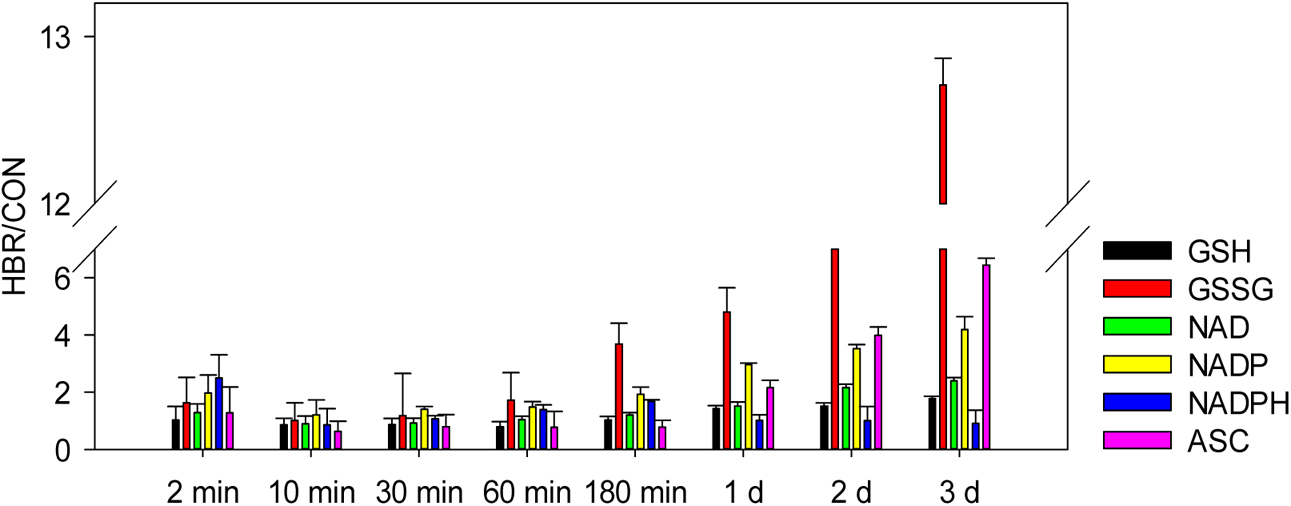
Redox-sensitive metabolites: Relative signal intensity after exposure to high light (HBR) versus normal light (CON). Structural identifiers for metabolite abbreviations are given in Table-SI 5.

In this context, malate valves act as powerful systems for balancing redox conditions within and between subcellular compartments in plant cells by exporting and accumulating excess reducing equivalents in the form of malate (Selinski & Scheibe, 2019). Isoforms of malate dehydrogenases (MDHs) are components of these valves and catalyze the reversible interconversion of malate and oxaloacetate. As shown in Figure 3 (gene group K), most MDH-encoding genes are transcriptionally activated under high light conditions within the scale of hours. Interestingly, at 72 h key genes encoding enzymes of the malate valve, such as the chloroplast-localized NAD-dependent (AT3G47520) and NADP-dependent (AT5G58330) MDH, returned to values comparable to those under control light conditions (Figure 3, Table-SI 4).

We next assayed the ratio between oxidized and reduced thiol groups in relevant proteins and found a tight maintenance of the redox ratio for all three measured time points (30 min, 180 min and 3 d) (*Table-SI 1*) (Nietzel et al, 2020). Together the results infer close maintenance of redox homeostasis under normal and also under high light, which is maintained by metabolic rearrangements as depicted above. In the course of continuous light stress this is accompanied by the accumulation of small redox molecules, which are seen as footprints of redox stress defense or can help quench ROS at enhanced cellular levels.

#### Metabolites of the MEP pathway are induced upon high light exposure

In view of its dual function as a precursor in isoprenoid metabolism and in retrograde signaling, we analyzed the time profile of methylerythritol 2,4-cyclodiphosphate (MEcPP) as well as other MEP pathway intermediates (Jiang et al, 2020; Xiao et al, 2012). Similar to other sugar phosphates and intermediates of photosynthetic dark reactions, all detectable MEP pathway intermediates showed increased levels within 2 minutes in light-stressed samples (Figure 4E). However, unlike for sugar phosphates and CBB cycle intermediates, we noted an amplified and prolonged increase for MEP pathway intermediates under light stress. The most prominent increase was observed for MEcPP and hydroxy-methyl-butenyl diphosphate (HMBPP) between 60 and 180 min in light-stressed plants with MEcPP levels at subsequent time points going down but still elevated levels and HMBPP remaining high relative to the normal light controls. By contrast, none of the corresponding MEP pathway genes shows a notable differential expression between 0-180 min when comparing high and normal light, indicating that the regulation of the MEP pathway operates at the post-translational level during this early phase (Figure 8A, *Table-SI 4*). Moreover, in plants, the conversion of MEcPP and HMBPP is mediated by reductases that use plastidic ferredoxins as electron donors (Seemann et al, 2006) (Figure-SI 1). Therefore, the flux through the MEP pathway also depends on the availability and redox state of ferredoxins. In support of increased production of plastidic ferredoxins, relative transcription for ferredoxin 1 of PS1 (FD1) (AT1G10960) constantly increased between 2 min and 24 h consistent with the accumulation of MEcPP and HMBPP over time under high light (Figure 3, gene group D, *Table-SI 4*, Figure 4E). The redox state of thiol groups in FD1 after 30 and 180 min reflected a slight but not significant shift towards the formation of more oxidized disulfides in light stressed plants (*Table-SI 1*). Moderate differences in gene expression from the MEP pathway under high light conditions were observed only at later time points, consistent with the increased accumulation of plastidial isoprenoids (see below).

**Figure 8.**
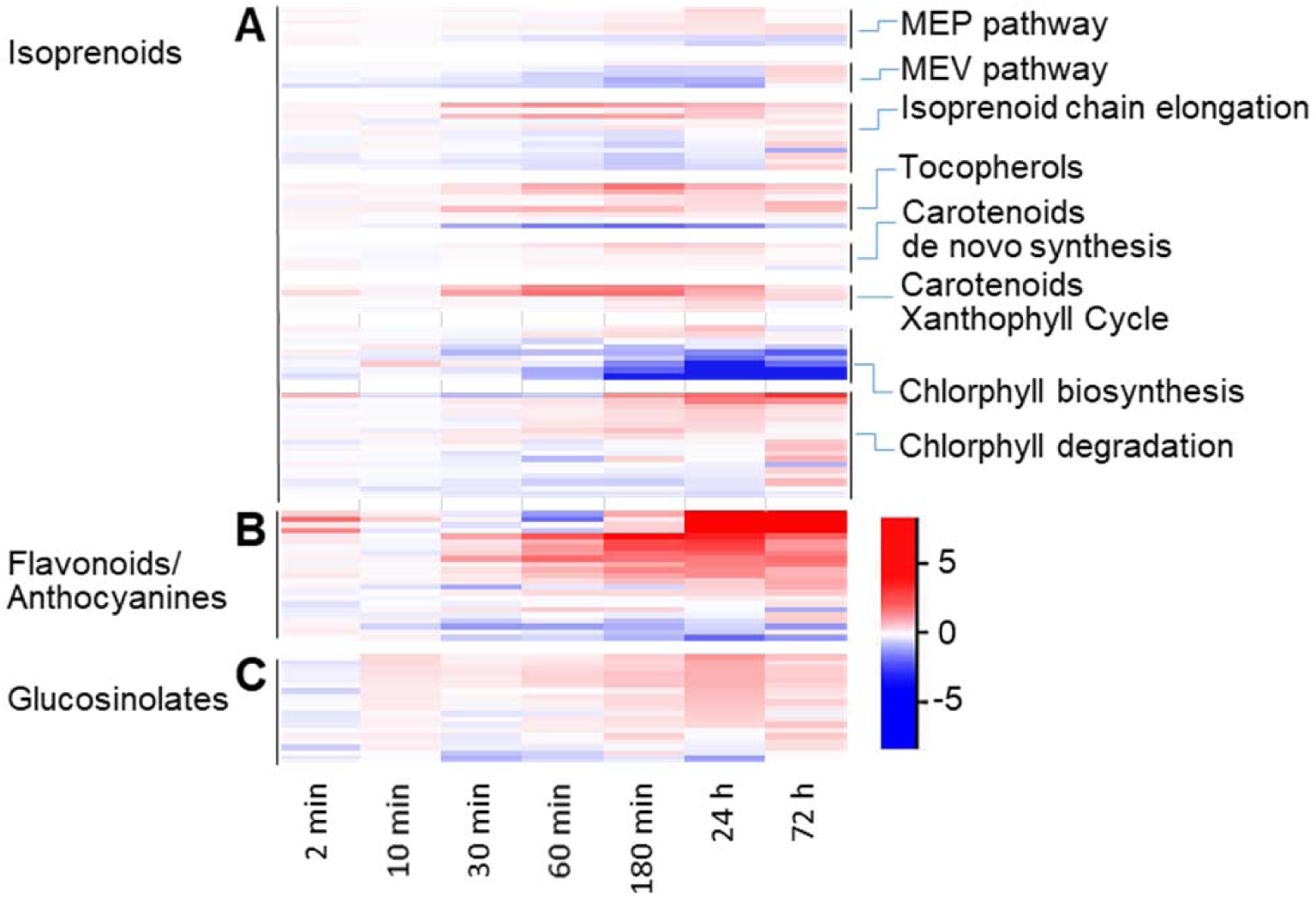
Differential gene expression profile within selected groups of transcripts involved in the formation of pigments, light-harvesting and light-protective metabolites. **A**, Isoprenoids. **B**, Flavonoids/anthocyanins. **C**, Glucosinolates. The graph depicts log_2_-fold changes between high light and control light (n=3) at a given sampling time (all data are unscaled).

Precursors of isoprenoids in plants can also be supplied by the cytosolic mevalonate (MEV) pathway. DEGs of biosynthetic enzymes of the MEV pathway revealed a downregulation with increased light intensity, which opposes the trend seen for the MEP pathway (Figure 8A). Intermediates of the MEV pathway were too low to be detected reliably, suggesting this pathway does not play a significant role in the acclimation to high light. Under high light, a similarly rapid increase in the FIE was observed for MEcPP and DMAPP. Interestingly, under normal light conditions, DMAPP was more quickly replenished with ^12^C than MEcPP and the isotopic distribution of both metabolites was comparable (Figure 5, Figure-SI 3). For instance, after 180 min the N+0 isotopolog contributes to more than 76% to the entire pool size of DMAPP while for MEcPP it is only 48%. This could indicate the presence of a second pool of DMAPP that does not derive from MEcPP (Figure-SI 1), or that there are two pools of MEcPP, of which one is replenished faster than the other. Under light stress the N+0 isotopolog for both intermediates of the isoprenoid biosynthesis increased more quickly than in normal light with maximum N+0 accumulation at around 180 min. The fact that MEcPP and its downstream product DMAPP show the same profile for FIE supports a dominant contribution of the biosynthetic flux into isoprenoids via the plastidic MEP pathway in excess light. The flux into MEcPP and DMAPP is equally rapid as observed for most sugar phosphates.

#### High light induces the sequential accumulation of specialized metabolites

Using MetFamily, we were able to annotate metabolite families from untargeted LC-QToF-MS/MS runs that showed increasing accumulation under light stress (Treutler et al, 2016). Those include flavonoids/anthocyanins, glucosinolates, arabidopsides, sulfoquinovosyl diacylglycerols (SQDG), and β-sinapoylglucose (m/z 385.12;4.7 min) within the clade of hydroxycinnamic acid derivatives (Figure-SI 7). For illustration, we selected representative substances from those families of secondary metabolism (Figure 9). Interestingly, individual compound classes show distinct and sequential accumulation over time. For instance, phenylpropanoids, anthocyanins, tocopherols, and stress-responsive glucosinolates accumulate already after 24 h consistent with the transcriptional induction of the corresponding biosynthetic genes preceding the accumulation of these metabolites (Figure 2C cluster 6, Figure-SI 2, Figure-SI 2, Figure 8A-C). By contrast, carotenoids and their ring-oxidized versions (xanthophylls) did not increase until after 48 h of light stress (Figure 9A-F). Likewise, plastoquinone-9, which is involved in the electron transfer between PSI and PSII, was found to be enhanced under light stress only after 48 h (Figure 9G). Then again, chlorophyll levels did not change throughout the experiment, despite transcriptional downregulation of several genes, which are involved in chlorophyll biosynthesis (Figure 9H, Figure 8A, Table-SI 4). Chlorophyll biosynthesis is negatively regulated by early light-induced proteins (ELIPs), which is in accordance with the early transcript upregulation of ELIPs observed under high light (Figure 3C, Table-SI 4) (Tzvetkova-Chevolleau et al, 2007). Simultaneously, several transcripts encoding chlorophyll degradation were upregulated during the late phase under high light irradiation (Figure 8A).

**Figure 9.**
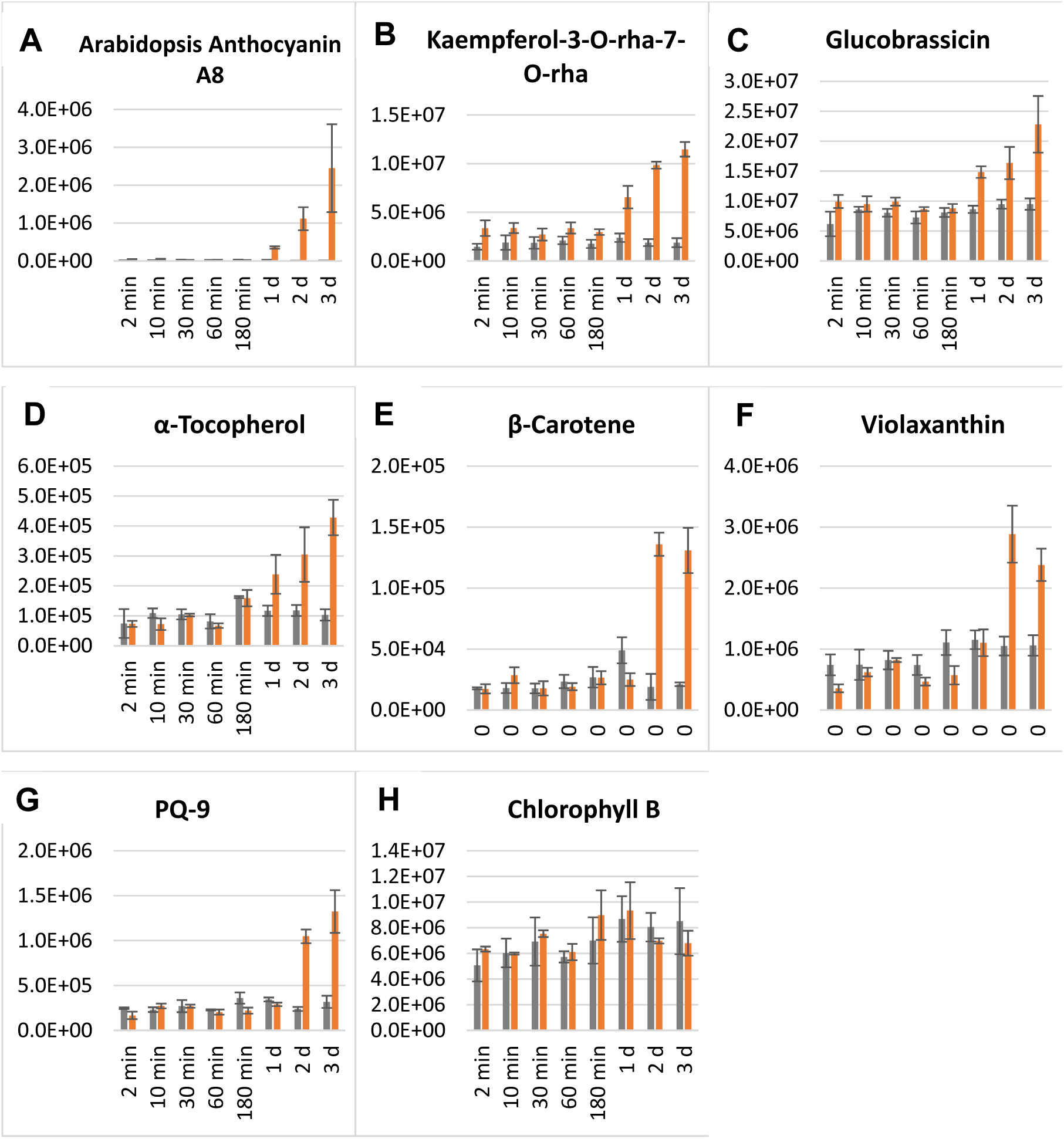
Time profile of selected photoprotectants and small molecules involved in light harvest. **A**, Anthocyanins. **B**, Flavonoids. **C**, Glucosinolates. **D**, Tocopherols. **E**, Carotenoids. **F**, Xanthophylls. **G**, Plastoquinones. **H**, Chlorophylls. All data are presented as peak area ± standard deviation (n=3).

Taken together, we show that metabolic acclimation to light stress proceeds in phases where photoprotectants need to be produced first. Since biosynthesis of large amounts of these compounds is time consuming significant accumulation is seen not before 24 h. Corroborated by the observation of transcriptional downregulation of light harvesting complexes during the first 24 h de novo carotenoid biosynthesis is also halted. Despite transcriptional downregulation of PS, LHC persists until 72 h carotenoid biosynthesis is activated by continuous light stress, which causes carotenoid accumulation after 2 d (Figure 3 panel B, Figure 8 panel A, Figure 9).

## Discussion

### Under Light Stress Redox Homeostasis is Maintained by Three Key Responses

In oxygenic photosynthesis, light produces ATP and NADPH in the plastids via linear electron flow. Yet, in plant leaves, continuous transfer of electrons through both photosystems by intense light irradiation can induce stroma over-reduction through permanent formation of reduced thioredoxins, ferredoxins, and NADPH (Endo et al, 2005). All of these compounds feature very low mid-point reduction potentials and therefore can act as strong reducing agents (Kozuleva & Ivanov, 2016; Scheibe & Dietz, 2012; Vogelsang & Dietz, 2020). Thus, redox shifts to a more reduced state, particularly when combined with an increase of these strong reductants in plant leaves would promote a partial reduction of molecular oxygen and ROS formation (Asada, 2000). On the other hand, insufficient supply of reducing power would impair the biosynthesis of essential compounds such as isoprenoids or lipids. Therefore, the availability of NADPH, the strongest reductant in a photosynthetic cell, is delicately balanced as observed (Figure 7).

In agreement with Hashida *et al*. long-term illumination increases the pool of NADP+ (Hashida et al, 2018). Notably, in redox couples with strong reducing agents the corresponding redox partners possess only weak oxidation power. Therefore, slowly increasing levels of oxidized glutathione (GSSG), NADP and NAD and increasing ratios between GSSG/GSH or NADP/NADPH in light-stressed plants are footprints of ROS attenuation, e.g. by the glutathione-ascorbate cycle, but do not mirror large changes in the cellular redox state. Similarly, slowly increasing levels of reduced ascorbate contribute to acclimation to high light as a high ascorbate pool will increase the capacity to recycle GSH and NADPH but store reducing equivalents in the form of a mild reductant ((Matsui et al, 2015), Figure-SI 4). Tight redox homeostasis is also reflected by nearly constant ratios between oxidized and reduced thiol groups in selected proteins involved in photosynthesis (Table-SI 1). This differs from findings when studying night to day transitions, where the redox state is significantly altered within minutes (Zimmer et al, 2021).

#### Under light stress metabolic pathways that produce NADPH are downregulated

Thus, reduced light harvesting (*de novo* chlorophyll biosynthesis, LHC) combined with the upregulation of cyclic electron flow and PTOX constitute a set of primary defense reactions to high light stress in order to avoid chloroplast over-reduction (Figure 8A, Figure 3B,E,F). Increased cyclic electron flow maintains high pH gradients at the thylakoid membrane for ATP production but limits the photosynthetic resupply of NADPH by PSI and the oxygen production in PSII (Joliot & Johnson, 2011). Besides the photosynthetic linear electron flow NADPH can also be supplied by alternative routes such as the oxidative pentose phosphate pathway, via malic enzymes and by action of isocitrate dehydrogenases. Transcripts of corresponding key enzymes were all downregulated by high light suggesting contributions in all cell compartments showing concerted reactions in order to avoid over-reduction (Table-SI 4).

#### Reactions that consume reducing equivalents are upregulated under light stress

When exposed to high light, plants increase NADPH-consuming CO_2_ assimilation via the CBB cycle and flux through the MEP pathway, which is observed by accelerated label incorporation into metabolic intermediates and indirectly by increased starch and isoprenoid accumulation relative to control light (Figure 3G, Figure 6, Figure 9). CBB cycle and photorespiration are intrinsically tied together by the oxygenase activity of Rubisco, leading to transient accumulation of the toxic intermediate 2-phosphoglycolate (Figure 3H, Figure 5, Figure-SI 3)(Nunes-Nesi et al, 2010). Detoxification of 2-phosphoglycolate by photorespiration involves four cell compartments and finally leads to production of NADH in the mitochondria, where it is used to drive oxidative phosphorylation (Figure 10). Peroxisomal hydroxypyruvate reductase (HPR), as a central part of 3-PGA salvage in the photorespiratory pathway, produces NAD(P)H, which then supports reduction of oxaloacetate to malate (see below). Thus, photorespiration redistributes reducing power between subcellular compartments (Fernie & Morgan, 2013; Lim et al, 2020; Nunes-Nesi et al, 2010; Walker et al, 2020), a process that is enhanced in the context of high light stress.

**Figure 10.**
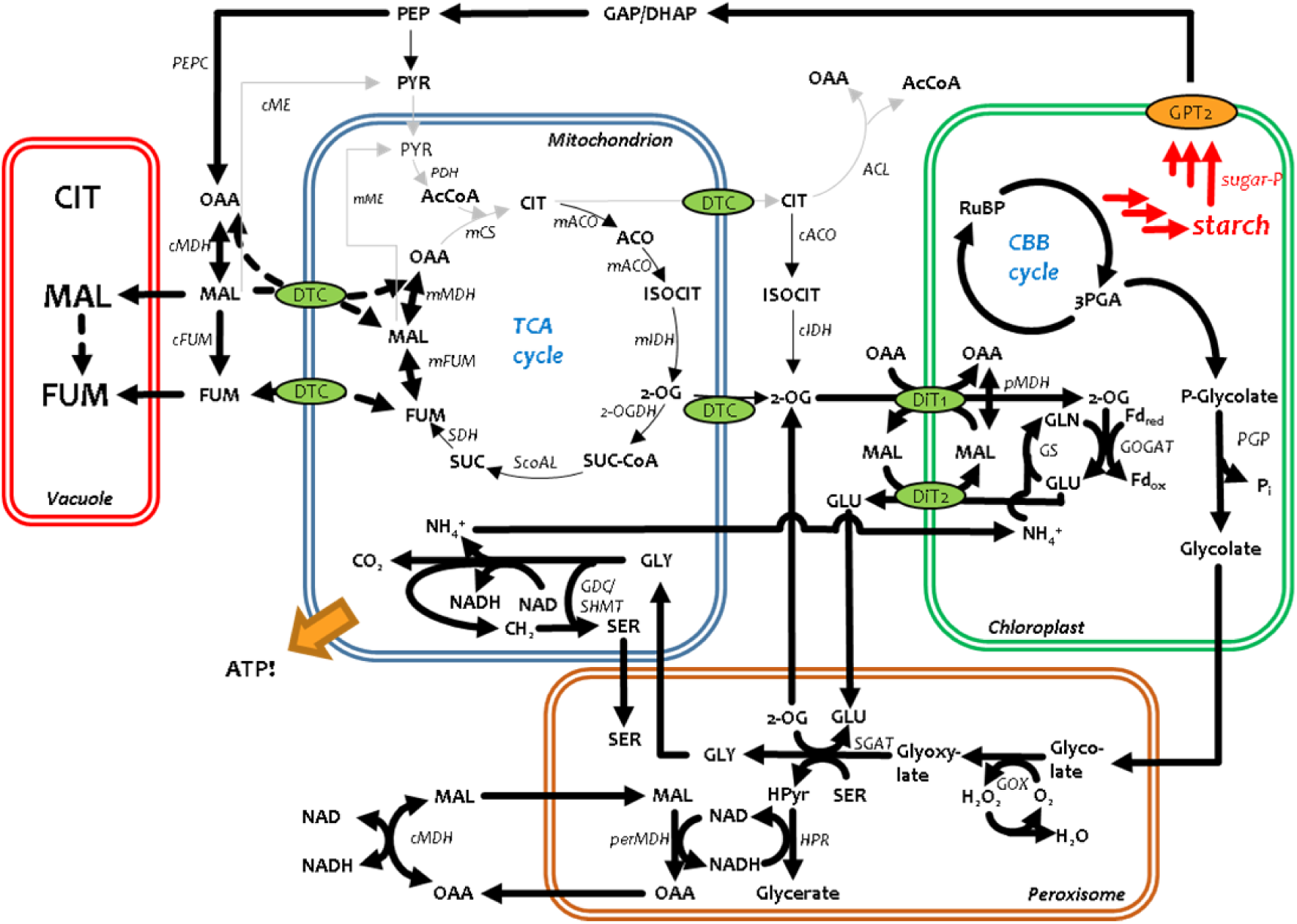
Central C and N metabolism under light conditions. Bold arrows symbolize fluxes that are enhanced under high light; narrow grey arrows represent fluxes that are decreased under high light.

Transcriptional activation of PTOX, which catalyze the reduction of molecular oxygen using reduced plastoquinones as substrates, indicate that PTOX functions as an essential outlet in photosynthetic electron transport of light-stressed vascular plants as described for photosynthetic microalgae (Curien et al, 2016; Saroussi et al, 2019). Besides PTOX, reduction of O_2_ on the stromal side of thylakoid membranes is induced in a light-dependent manner by the Mehler reaction (Curien et al, 2016). Hereby, cell-toxic ROS such as H_2_O_2_ are formed. Detoxification of these ROS is supported by transcriptional activation of genes involved in redox metabolism and confirmed by the increase in oxidized metabolites (GSSG, NAD^+^, NADP^+^).

#### Accumulation of C4 acids in vacuoles represents a storage form of reduced carbon under high light

During the day, biosynthesis of starch constitutes a major sink for photosynthetic reducing power. However, in chloroplasts under stress, excess reducing equivalents are believed to be redirected to other sinks (Selinski & Scheibe, 2019; Selinski & Scheibe, 2020). This form of cellular redox poise is promoted by the export of reducing equivalents from the chloroplast via the malate valve (Igamberdiev & Eprintsev, 2016; Selinski & Scheibe, 2019). As components of malate valves, isoforms of malate dehydrogenases catalyze the interconversion of malate and oxaloacetate in chloroplasts, the cytosol, peroxisomes, and mitochondria (Dao et al, 2021). *Arabidopsis thaliana* possesses nine genes encoding MDH isoenzymes of which seven showed significant upregulation under light stress (Table-SI 4). Correspondingly, our data further show strong diurnal *de novo* synthesis of malate and fumarate under light stress resulting in important accumulation of both acids (Figure-SI 8, Figure 5). Absolute quantitation after non-aqueous phase separation showed large pool sizes and vacuolar localization for citrate, malate, and fumarate in Arabidopsis leaf (Szecowka et al, 2013). Hence, we can conclude that increased levels of malate and fumarate as observed under light stress contribute to a second subcellular pool of “reduced carbon” which is localized primarily in the vacuoles. Surprisingly, on a molar scale, this pool is much bigger than the carbon stored as starch, indicating the pivotal role of MDHs when light is excessive.

Taken together, permanently light-stressed plants apparently adopt three strategies in order to maintain redox homoeostasis. One is to reduce further production of strong reductants. The others are to attenuate excess reducing power by activating processes that consume reducing equivalents or distribute them across the cell to other subcellular compartments for storage and later consumption.

### Photorespiration and malate metabolism jointly fuel mitochondrial activity

#### Forward flux through the TCA cycle is suppressed by high light while C4 acids are produced via anaplerotic routes

In Arabidopsis under light stress, malate and fumarate not only accumulated over time but were synthesized *de novo* at higher rates than in low light. However, synthesis of malate and fumarate via the clockwise action of the TCA cycle is negligible because virtually no isotope label enrichment was found in citrate and *cis*-aconitate over 300 min of light illumination, raising the question about the origin of these C4 acids. Malate can also derive from anaplerotic reactions of PEP and CO_2_ via phosphoenolpyruvate carboxylase (PEPC) and reduction of oxaloacetate (Figure 10). PEPC was shown to be diurnally activated by post-translational modification, which would be in accordance with the absence of transcriptional activation of PEPC under high light (Asai et al, 2000; Uhrig et al, 2019) (Table-SI 4). Previous work on illuminated *Xanthium* leaves indicated that under conditions of active photosynthesis the TCA cycle can supply citrate for the synthesis of cytosolic acetyl CoA, 2-oxoglutarate and glutamate (citrate valve) while accumulation of malate regulates the redox balance in different cell compartments via the malate valve (Igamberdiev & Eprintsev, 2016; Tcherkez et al, 2009). Earlier, positional forward labeling by feeding 1-^13^C- and 2-^13^C-pyruvate inferred stronger anaplerotic formation of malate when atmospheric CO_2_ levels were depleted (Tcherkez et al, 2008). Notably, conditions of low CO_2_/O_2_ favor the photorespiratory flux, which in turn correlates with increased anaplerosis. A similar situation may arise under light stress where the CO_2_/O_2_ ratio in the leaf decreases due to massive oxygen production and stomatal closure (Devireddy et al, 2018). In contrast to pyruvate-feeding studies conducted by Tcherkez and coworkers, we found very little turnover in citrate, aconitate, 2-oxoglutarate and glutamate during the first 300 min, showing that in Arabidopsis de novo C5-body formation does not operate in the light. By contrast, since in our chase experiment PEP was rapidly enriched with ^12^C-carbon, we conclude that under light stress fresh malate and fumarate are almost exclusively produced via anaplerosis. Moreover, the pyruvate dehydrogenase complex (mPDC) is inactivated by light, an effect that is enhanced by photorespiration (phosphorylation) (Gemel & Randall, 1992). This would explain the absence of label in citrate and downstream metabolites. Artificial pyruvate substrate feeding, as performed by Tcherkez and collaborators may lead to high intracellular pyruvate levels that are not physiological and thereby activate PDC (Tovar-Méndez et al, 2003). This may then lead to citrate, 2-oxoglutarate and glutamate labeling, which we and others did not observe when ambient conditions prevailed (Figure 5, Figure-SI 3) (Gauthier et al, 2010).

#### Mitochondrial ATP generation under light stress is fueled by glycine oxidation

Interestingly, transcripts encoding enzymes of the mitochondrial electron transport chain were quickly upregulated under high light inferring enhanced mitochondrial activity (Figure 3M, Table-SI 4). Under light stress, we noted a faster incorporation of ^12^C in several photorespiratory intermediates, indicating an enhanced flux through photorespiration. This increased flux under high light would lead to enhanced glycine oxidation by the glycine decarboxylase complex (GDC) in leaf mitochondria, which can provide the NADH required to drive oxidative phosphorylation in the absence of a fully cyclic TCA cycle (see above) (Figure-SI 8, Figure-SI 3). Moreover, GDC releases ammonium, which also provides an important link to nitrogen metabolism (Figure 10). The low k_cat_ values of GDC, (in the order of 10 s^−1^) (Bykova et al, 2014), one of the most abundant proteins in planta, would explain the transient accumulation which has been shown to enhance the inactivation of mPDC in vitro by stimulating the pyruvate dehydrogenase kinase that catalyzes the inactivating phosphorylation (inactivation) of the mPDC (Schuller & Randall, 1989). Thus, increased photorespiration leads to further inhibition of the oxidation of pyruvate and by the same token prevents further production of reducing equivalents via the TCA in the mitochondria.

Further adjustments in the redox balance in the mitochondria can be achieved through the activity of malate dehydrogenase (MDH). The upregulation of transcripts for mMDH1 and mMDH2 supports this hypothesis.

Taken together, our data support the conclusion that under light stress malate metabolism and photorespiration are tightly orchestrated with nitrogen metabolism. Photorespiratory glycine oxidation probably strongly energizes mitochondrial membranes to maintain ATP homeostasis in absence of a full TCA cycle (Figure 10). Under high light, salvage of oxaloacetate to support a vital export of reducing equivalents from chloroplasts in form of malate is achieved by activation of PEPC and several MDH.

#### Nocturnal metabolism reactivates forward flux over TCA cycle and C5-body renewal of glutamate

Strongly increased ^12^C-enrichment in citrate and *cis*-aconitate was observed only in days 2 and 3, i.e. after a period of darkness, indicating that citrate synthesis occurs during the night. Incorporation of ^12^C into citrate is likely to come from the glycolytic decomposition of non-labeled glucose originating from starch, which was formed during the previous day. The vacuolar pool of malate and fumarate may also replenish the TCA cycle either directly after dicarboxylate transport into the mitochondria or by nocturnal activation of malic enzyme (Badia et al, 2015; Tronconi et al, 2008) (Figure-SI 9). Thus, isotopic patterns observed after the first night imply that the TCA cycle must have been active in clockwise direction during the night. Transcriptional upregulation during the night or extended night was also reported for cytosolic isocitrate dehydrogenase (cICDH) (At1g65930) and glutamate dehydrogenases (GDH) (At5g07440, At5g18170) and GDH showed increased nocturnal activity (Gibon et al, 2006). Collectively, and since large parts of N+0 had formed after one night (day 1), *de novo* synthesis of the C5 body in glutamate must have largely occurred during the night. The extent of this nocturnal de novo synthesis was significantly higher when the plants were exposed to light stress during the preceding day (cf. 2-OG and GLU in Figure-SI 3). Ammonium represents an end product of photorespiration, which is consumed by GOGAT and glutamine synthase (Figure 10). Apparently, GOGAT and glutamine synthase intensively recycle the same C5 body in the light but extensively renew the C5 pool during the night (Figure-SI 9, Figure-SI 3).

### Concerted Responses to High Light Stress

Overall, our data illustrate well-orchestrated responses of the Arabidopsis model towards high light irradiation, which can be classified into three time layers: perception (few seconds to minutes), early responses (minutes to hours), and acclimation (hours to days) (Figure 1).

#### Rapid Activation of Heat Shock Proteins Indicates the Importance of Chaperone Functions Under High Light

There are only few transcripts that are upregulated within minutes upon high light exposure. Among those are several transcripts encoding heat shock proteins (HSP) and their regulating factors (HSF) showing that stabilizing proteins to ensure correct folding or helping to refold proteins that were damaged by high light requires an immediate response. Previous studies point to a relationship between HSP expression and light (Dickinson et al, 2018). By contrast, Huang et al. claim that by cooling down light-stressed plants, they could mostly eliminate the induction of HSP (Huang et al, 2019). However, their experiments were done in constant light, which was shown to eliminate the induction of HSF observed during a regular day-night cycle (Dickinson et al, 2018). Therefore, although the slight temperature increase we observe (2.5°C) cannot be fully neglected it appears that in adult plants high light itself rapidly activates HSP. Ongoing transcriptional upregulation of HSP throughout further time points indicates that heat shock proteins remain an important factor in the acclimation to long-term light stress.

Stable isotope labeling, transcriptional and metabolic time profiles in this study further revealed that increased carbon fixation, stress avoidance and stress attenuation reactions are simultaneously activated within the scale of tens of minutes in response to high light. In contrast to most transcripts, metabolite levels of central carbon metabolites can change instantly. For instance, transcriptional activation of genes of the CBB cycle and photorespiration occurred only after 30-60 min but the levels of several metabolic intermediates and incorporation of fresh carbon increased already after 2 min under high relative to control light (Table-SI 4, Figure 4A and B, Figure 5, Figure-SI 3). In accordance with our current state of knowledge, elevated flux through both pathways at a very early stage of light stress is not controlled by transcriptional activation (Michelet et al, 2013). However, significant changes in the redox state of thiol groups, when comparing high and normal light conditions were not observed either. This suggests additional post-transcriptional modes of regulation. Phosphorylation of Rubisco-activase for example plays a role in the regulation of its activity and thereby can rapidly impact Rubisco activity (Boex-Fontvieille et al, 2014; Kim et al, 2019).

Many transcriptional and metabolic regulations ensure redox homeostasis but also reflect conditions of long-lasting modifications of the photosystem apparatus, which for instance manifest in strong increases of carotenoid and plastoquinone pools until day 2 and 3. In the course of such acclimation to high light many of early regulated gene families show only little or no differential regulation anymore towards the last sampling point after 3 days. Those families reflecting these acclimation processes include genes encoding light harvesting complexes (LHC), chlorophyll biosynthesis, cyclic electron flow, PTOX and NADPH-forming non-photosynthetic responses (Figure 3).

The metabolic levels of light harvesting chlorophylls remained unaffected despite the activation of chlorophyll degradation during the last two time points (Figure 3B, Figure 8A, Figure 9H, Table-SI 4). This is explained by high chlorophyll levels that were present already before the experiment began and by a slow turnover of chlorophylls.

#### Sequential accumulation of light protective and photosynthesis-supporting metabolites

Applying mass spectral clustering techniques we annotated several metabolite families (Figure-SI 6, Figure-SI 7))(Treutler et al, 2016). Accumulation under high light conditions was exemplarily shown for individual members of those metabolite families known to be involved in the response to light stress (Figure 9) (Kobayashi, 2018; Kumar et al, 2020; Neugart et al, 2019; Tohge et al, 2016; Zhang et al, 2018). However, our data show that biosynthesis of individual metabolite families occurs in a sequential order. Phenylpropanoids and tocopherols began to accumulate much earlier than carotenoids and plastoquinone-9. The former are thought to function as photoprotectants in the absorption of energy-rich light and as ROS scavengers. The latter two are rather involved in light harvest and mediation of electron transport. Hence, we conclude that in the concert of acclimation, only after high light protecting metabolites accumulate, do photosynthetic processes return closer to levels seen before the light stress.

#### The MEP pathway has a role in the early response to high light

Unlike other sugar phosphates, MEcPP displayed a distinct accumulation already within minutes with a peak at 3 hrs. This increase coincided with the induction of stress-responsive genes such as fatty acid hydroxyperoxide lyase (HPL, AT4G15440) that are known to be induced by the accumulation of MEcPP (Figure 4, Figure 3L and Table-SI 4). We also observed increasing levels of HMBPP, which is the direct product of HDS (4-hydroxy-3-methylbut-2-enyl diphosphate synthase), the enzyme whose substrate is MEcPP (Figure-SI 1) (Wang et al, 2020; Xiao et al, 2012). This indicates an increased flux through the MEP pathway upon high light stress. A similar isotopic distribution of MEcPP and its downstream product DMAPP during high light irradiation indicates that the MEP pathway is the predominant synthetic route of isoprenoids. By contrast, under normal light we observed faster reverse labeling of DMAPP than MEcPP than under high light and a relative upregulation of genes encoding the MEV pathway. This could indicate an increased contribution of the MEV pathway to the synthesis of DMAPP with lower light irradiation (Figure-SI 3). However, the metabolite levels of the MEV pathway are so low (most of them are not detectable) that a significant contribution to the global pool of IPP/DMAPP is unlikely. Alternatively, the contribution from a second pool of MEcPP and its derivatives could also explain the different labeling pattern between MEcPP and DMAPP (Gonzalez-Cabanelas et al, 2015).

A notable differential expression of MEP pathway-encoding genes was not observed during the first hours of light stress indicating flux control through the MEP pathway via post-transcriptional mechanisms. Noteworthy, the two enzymes that use MEcPP and HMBPP as substrates – respectively HDS and HDR (4-hydroxy-3-methylbut-2-enyl diphosphate reductase) - contain iron-sulfur clusters (Seemann et al, 2005; Wang et al, 2020; Wolff et al, 2003). Additionally, the reduction step from MEcPP to HMBPP depends on the electron transfer from reduced ferredoxins (Seemann et al, 2006). Therefore, the flux through the MEP pathway should be highly dependent on the redox state in the chloroplasts. In Zymomonas, which exclusively produces isoprenoids via the MEP pathway, oxidative stress was induced by a shift from anaerobic to aerobic growth, causing temporal accumulation of both MEcPP and HMBPP (Martien et al, 2019). Similarly, high light should induce oxidative stress, with impact on the redox state of plastidic ferredoxins, HDS and HDR. The actual electron transfer occurs in iron-sulfur reaction centers transfer by changing the redox state of iron between Fe2+ and Fe3+. This transfer, however, could not be directly assessed. As a surrogate we analyzed the redox state of thiol groups in HDS, HDR and plastidic ferredoxins but to our surprise all were tightly redox balanced throughout the experiment. Thus, the key factors that regulate the flux through the MEP pathway remain elusive and should be addressed in future work.

The combination of transcriptome, metabolome, stable isotope labeling and redox-proteomics allowed to uncover multiple and sequential events when plants are exposed to light stress. Importantly, metabolic profiling and labeling unraveled processes, especially those happening within short times, that are otherwise not detected by transcriptomics and proteomics. During this concerted response, we observed in the early phases, a number of mechanisms that are activated to maintain the redox homeostasis across several cellular compartments, while at later time points, accumulation of light protective metabolites supports a remodeling of the photosynthetic apparatus.

## Materials and Methods

### Plant culture and harvest

#### Arabidopsis thaliana

Columbia was grown for seven weeks on soil under short day conditions (8h light period, 22 °C (130 µmol m^−2^ s^−1^, 16h dark period, 20 °C, 70% humidity). During this time plants form rosettes of ca. 50 mm diameter without flowering. Then, for 7 days, plants were moved to long day conditions with a light period of 16 h, the other conditions remaining unchanged. Light stress experiments were performed on 40 plants, while another 40 plants served as controls. For both, mobile LED-light panels were used (Rhenac GreenTech AG, Hennef, Germany). For stress conditions, a light intensity of 1330 µmol m^−2^ s^−1^ (230 nm – 1000 nm), and for control conditions a light intensity of 130 µmol m^−2^ s^−1^ was applied. Light spectra between 230 nm and 1000 nm were recorded by a Specbos 1211UV (JETI, Jena, Germany) (Figure 1A-B). In order to minimize temperature differences at the leaf surface, the distance between the LED source and the plant panel was set to 0.72 m. For each time point three individual rosettes were cut and immediately quenched in liquid nitrogen (<1s). For experiments without labeling, individual rosettes were separately extracted and measured. For pulse-chase labeling experiments, 3-4 rosettes per time point were pooled before extraction.

### Pulse-Chase-^13^C-Labeling

80 seven-week-old plants were cultivated as above. Thereafter, plants were put into an enclosure (Phytobox Labelbox, Elektrochemie Halle, GmbH) (Balcke et al, 2017). Therein plants were fumigated over 7 days with 450 ppm ^13^C-CO_2_ (99.3% isotopic purity, Euroiso-Top, Sait-Aubin Cedex, France,) under 16h long day light conditions of 130 µmol m^−2^ s^−1^ and 60% humidity. The controller of this enclosure was able to simultaneously detect ^12^C-CO_2_ and ^13^C-CO_2_ levels but could maintain constancy only for ^13^C-CO_2_ partial pressures. During the pulse ^13^C-labeling pre-phase plants continued to release ^12^C-CO_2_, which diluted the fractional enrichment of ^13^C-carbon in the metabolites. Particularly, carbon dioxide release by nocturnal plant respiration significantly contributed to an increase in the atmospheric share of ^12^C-CO_2_, which was manually removed by daily morning scrape-off of all carbon dioxide and replacement with fresh 500 ppm ^13^C-CO_2_. At the end of the pulse phase, when the cabinet was opened, 384 ppm ^13^C-CO_2_ and 125 ppm ^12^C-CO_2_, respectively, were detected.

### Metabolite extraction

Shock-frozen material was ground under liquid nitrogen conditions. 150 mg frozen fresh weight was balanced into wall-reinforced extraction tubes containing 1 steel bead (3 mm), 3 steel beeds (1 mm) and 200 mg of glass beads (0.75-1 mm) and maintained at temperatures ≤-80 °C throughout. For metabolite analysis, a modified 2-phase extraction was performed where 900 µL dichloromethane/ethanol (2:1, −80°C) were first added followed by 150 µL dilute HCl (pH 1.7). A first extraction was performed by cryo bead milling for 3*20 s (MP24, Biomedicals Inc., 4.0 m s^−1^). During this period the initial temperature increases from −80 °C to 14 °C. Following phase separation by centrifugation (2 min, 10,000 x g) the upper phase (containing hydrophilic metabolites) was collected and stored on ice. Then another 100 µL aqueous HCl was added, and the extraction and phase separation were repeated. Both aqueous extracts were combined and stored at −80 °C until analysis by methods 1 and 2. The remaining organic phase was collected and stored on ice, and the cell debris, which forms an interlayer between aqueous and organic phase, was re-extracted with 500 µL tetrahydrofuran (THF). Both organic extracts were combined, aliquoted and dried in a nitrogen stream. Aliquot 1, was dissolved in 80% methanol, centrifuged and analyzed by method 3. Aliquot 2 was dissolved in 2-propanol, centrifuged and analyzed by method 4.

### Transcriptomics

Shock-frozen material was ground under liquid nitrogen conditions. 100 mg of cryo-ground leaf powder was used to isolate total RNA using the RNAeasy kit from QIAGEN. The quality of total RNA was measured by evaluating of RNA integrity number (RIN) using BioAnalyzer quality control analysis (Agilent, Santa Clara, CA) and only samples with a RIN > 8 were used for cDNA synthesis. 1 μg of RNA treated with DNase (RQ1, Promega) was reverse-transcribed by Superscript III (Life Technologies). Strand specific cDNA library was subjected to Novaseq sequencing and produced 19.8-33.3 million reads (150 bp paired end) per sample (Data-SI 1). All data from 45 different treatments was analyzed using our bioinformatics pipeline in the platform usegalaxy.eu (Afgan et al, 2018). FastQC v 0.11.5 and MultiQC were used to assess the quality of reads (Andrews, 2010; Ewels et al, 2016). The reads for each sample were trimmed and filtered using Trimmomatic v0.32 (Bolger et al, 2014). High quality reads with a Phred score >30 were aligned to the Arabidopsis TAIR 10 genome using HISAT2 to generate SAM (Sequence Alignment/Map) (Kim et al, 2015). SAM files were used to count aligned reads using htseq-count and generate count tables, which were submitted to DESeq2 for determination of differentially expressed genes (DEG) (Anders et al, 2014; Love et al, 2014). A threshold of a log2 fold change ratio ≥ 1 was used to define significant DEG after cutting off the data at a false discovery rate (FDR) value ≤ 0.05 for all unigenes with ≥0.5TPM (Transcripts Per Kilobase Million). The raw data for RNA sequencing and sampling details are deposited at the NCBI as Bioproject PRJNA817005 (Data-SI 1). By use of MAPMAN 4.0 we further categorized all detectable transcripts and performed k-means time cluster analysis (Schwacke et al, 2019). Gene ontology (GO) enrichment analysis was conducted by DAVID V. 6.8 (Huang et al, 2009a; Huang et al, 2009b).

### Metabolite profiling

#### Method 1. Targeted analysis of hydrophilic compounds of the CCEM

5 uL of the aqueous phase were injected to an Acquity UPLC (Waters Inc.) equipped with a Nucleoshell RP18 column (Macherey & Nagel, 150mm x 2 mm x 2.1µm) using tributylammonium as ion pairing agent. Solvent A: 10 mM tributylamine (aqueous) acidified with glacial acetic acid to pH 6.2; solvent B acetonitrile. Gradient: 0-2 min: 2% B, 2-18 min 2-36% B, 18-21 min 36-95% B, 21-22.5 min 95% B, 22.51-24 min 2% B., The column flow was 0.4 mL min^−1^ throughout. The column temperature was 40 °C. Scheduled multiple reaction monitoring (MRM)-based metabolite detection was performed in negative mode electrospray ionization (ESI) on a QTrap6500 (AB-Sciex GmbH, Darmstadt, Germany: ion source gas 1: 60 psi, ion source gas 2: 70 psi, curtain gas: 35 psi, temperature: 450 °C, Ion spray voltage floating: and −4500V). MRM transitions were of 189 metabolites were previously signal-optimized and retention times determined (***Table-SI 1***).

#### Method 2A and 2B. Untargeted analysis of hydrophilic metabolites

5 uL of the aqueous phase were injected to an Acquity UPLC (Waters Inc.) equipped with a Nucleoshell RP18 column (Macherey & Nagel, 150mm x 2 mm x 2.1µm) applying LC-Method 1 with QToF-MS/MS readout (Method 2A, negative ESI) and regular RP18 UPLC (Method 2B for positive ESI). Solvent A: 0.3 mM ammonium formate (aqueous) acidified with formic acid to pH 3; solvent B: acetonitrile. Gradient: 0-2 min: 5% B, 2-19 min: 5-95% B, 19-21 min: 95% B, 21.1-24 min: 5% B. The column flow was 0.4 mL*min^−1^ throughout. The column temperature was 40 °C. UPLC-separated signals were analyzed by SWATH-QToF-MS/MS (TripleToF5600, AB-Sciex GmbH, Darmstadt, Germany) with negative (Method 1) positive electrospray ionization (Method 2) using these source parameters; ion source gas 1 60 psi, ion source gas 2 70 psi, curtain gas 35 psi, temperature 600 °C, ion spray voltage floating −4500 V/5500 V. The MS scanning parameters are given in Data-SI 2.

#### Method 3A and 3B. Separation of medium polar secondary metabolites

5 uL of aliquot 1 of the organic phase were analyzed as in Method 2. Here, for untargeted metabolite profiling negative (3A) and positive (3B) ESI was used.

#### Method 4. Separation of APOLAR metabolites

10 uL of aliquote 2 of the organic phase were injected to an Acquity UPLC (Waters Inc.) equipped with a Nucleoshell RP18 column (Macherey & Nagel, 150mm x 2 mm x 2.1µm) using a lipidomics gradient of solvent A: 10 mM ammonium acetate (aqueous) and solvent B: 60% acetonitrile/ 40% 2-propanol. Gradient: 0-1 min: 60% B, 1-8 min 60-87.5% B, 8-16 min 87.5-98% B, 16-20.5 min 98% B, 20.5-22 min 98-60 %B, 22-24 min 60% B. The column flow was 0.4 mL min^−1^ throughout. The column temperature was 45 °C. Positive electrospray ionization and SWATH-QToF-MS/MS was used as in Method 3.

#### Metabolomics data analysis workflow

All data obtained from untargeted metabolomics was analyzed using the workflow illustrated in Figure-SI 5. In brief, vendor raw data from Sciex (LC-MS) was first converted to analysis base file (*.abf) using the free Reifycs software (download: https://www.reifycs.com/AbfConverter/). Abf data was uploaded to MS-Dial ((Tsugawa et al, 2015), http://prime.psc.riken.jp/compms/msdial/main.html). MS-Dial performed peak picking, peak deconvolution, metabolite identification and chromatographic alignment of independent samples in the respective batch using the parameters in Data-SI 2Exported MS^1^ raw data (peak heights) and a msp-formatted MS/MS library of detected spectra from MS-Dial were imported to MetFamily ((Treutler et al, 2016), https://msbi.ipb-halle.de/MetFamily/) and processed with the parameters in Data-SI 2. Prenol lipids, tetrapyrrole pigments and metabolites of the specialized plant metabolism of *Arabidopsis thaliana* were assigned by authentic standards and using MetFamily, which allows the annotation of entire metabolite clusters based on the similarity of measured MS/MS spectra (Data-SI 2, Figure-SI 5)(Treutler et al, 2016).

#### Quantification of selected metabolites and isotopolog signals

Selected metabolites and isotopologs of labeled compounds were detected by ToF mass spectrometry and were quantified as area by Multiquant TF (Sciex). Using these methods, signals for oxaloacetate and succinate were too low or ambiguous. Since citrate and isocitrate have isobaric [M-H] molecular ions of 191.0197 Da and showed nearly co-eluting peaks our LC-TOF method was unable to discriminate between both acids. Because in Arabidopsis leaves, absolute quantification of acids showed that the pool size of isocitrate is only 1.7% of that measurable for citrate (Szecowka et al, 2013), the entire signal was fully attributed to citrate.

We here use fractional isotope enrichment (*FIE*), where the *FIE* determines the molar fraction of ^12^C atoms in the pool of ^12^C- and ^13^C-carbon atoms of a metabolite.

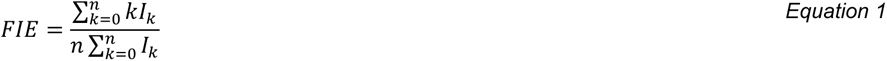

*with k ɛ (0..n) is the number of ^12^C isotopes and Ik the signal of the corresponding isotopologs with k isotopes. N is the total number of carbon atoms in a metabolite. MS^1^ isotopolog signals of [M-H] precursor ions were identified by high resolution mass spectrometry. Increasing FIE represent fluxes of ^12^C-carbon from ambient carbon dioxide into a metabolite’s pool, which was evaluated for control and light stressed plants*.

#### Quantification of malate, fumarate and citrate

20 to 60 mg homogenized fresh weight were extracted as described above, except three rounds of extraction were carried out, replacing the aqueous buffer three times. The combined aqueous phase gave a total volume of 450 mL of which 5 uL were measured by Method 1 and compared against freshly prepared authentic standards (Merck, Darmstadt, Germany). We wish to note that our acid quantification generated much higher absolute values than an earlier study (Szecowka et al, 2013). However, Szecowka et al. obtained their quantitative results from non-aqueous phase extraction, while we used a simple two-phase extraction approach, which we found to be exhaustive during previous method development.

### REDOX-Proteomics

Sample preparation and LC-MS/MS-based quantification of the cysteine oxidation status was performed as previously described in detail unless stated otherwise (Giese et al, 2022). 50 mg of frozen and finely ground leaf material was used per sample. Each labeling step was performed with 200 µg iodoTMTsixplex (LOT: VB291350, Thermo Scientific). Multiplex labeling reagents were swapped in between replicates. Samples were pooled as described in Giese et al. (2022) followed by fractionation on SDB-RPS stagetips as described in detail in (Lassowskat et al, 2017).

For LC-MS/MS analyses, 500 ng TMT-labelled peptide sample was measured on an EASY-nLC 1200 system coupled to an Orbitrap Exploris 480 mass spectrometer (Thermo Scientific, www.thermofisher.com). Liquid chromatography was performed using 20 cm fritless silica emitters with 0.75 µm inner diameter (New Objective, www.newobjective.com) containing reversed-phase 1.9 µm ReproSil-Pur C_18_-AQ (Dr. Maisch GmbH, www.newobjective.com). Peptides were separated on a 115-minute gradient of 4 % to 78 % acetonitrile with 0.1 % formic acid. Further details on chromatography and acquisition can be found in Table-SI 6.

Evaluation of MS raw data was performed with MaxQuant software (v 1.6.17.0) using Araport11 (v2016/06, www.bar.utoronto.ca/thalemine) as database and a reverse decoy database. Potential contaminants were included in the search. MS^2^-based quantification was performed by selecting the iodoTMT 6-plex preset with addition of the lot-specific isotope correction factors (LOT: VB291350). The default setting of carbamidomethylation was removed from fixed modifications. The ‘match between runs’ feature was enabled. Trypsin was set as protease and a maximum of two missed cleavages was allowed. Data analysis was performed using Perseus software (v 1.6.5.0, www.maxquant.org). Calculation of percentual oxidation status was performed as described in (Giese et al, 2022).

### Starch Quantification

Alcohol-insoluble residue (AIR) was obtained by sequentially washing 10 mg dry weight of ground leaf material once with 1 mL of 70% ethanol, once with 1 mL of chloroform:methanol (1:1 v/v), and once with 1 mL of acetone. After each washing step, the AIR was pelleted by centrifugation (20000g for 5 min), and the supernatant was carefully discarded. After the final wash, the pellet was resuspended in 300 µL of acetone, and dried in a speed-vac and balanced. Next, the pellet was resuspended in 400 uL 0.1 M NaOH and saponified at 120 °C for 30 min. After cooling down, the samples were neutralized by addition of 80 µL of an acetate buffer (0.5M HCl + 0.1 M HOAc 1:1, 4°C). The samples were resuspended and a homogeneous aliquot of 40 uL was hydrolyzed at 37 °C overnight using a mix 110 uL of µL amyloglucosidase (from *Aspergillus niger*, Sigma, 10102857001) + α-amylase (from pig pancreas, Sigma 10102814001) in an acetate buffer (pH 4.9). After digestion, glucose was determined by the D-Glucose Assay Kit (GOPOD Format, Megazyme) using D-glucose standards as reference.

## Acknowledgements

We would like to acknowledge the technical assistance of Anja Scherr-Henning, Mandy Dorn, Marvin Hempel and Paulina Heinkow (MSPUB, University of Muenster). We further would like to thank Stephanie Arrivault, MPI of Molecular Plant Physiology, Potsdam, Germany, Catalin Voiniciuc and Bo Yang (IPB) for help with the quantification of starch. This work was supported financially by the DFG (Deutsche Forschungsgemeinschaft) by grant number 469950637 to IF for proteomic analyses.

## Author contributions

GUB and AT designed the research, KV isolated RNA and performed statistical data analysis, JG and IF performed proteomics analysis, GUB conducted the light stress and labeling experiments and performed the metabolomics measurements and data analysis, GUB and AT wrote the article with contributions from the co-authors.

## Extended View

**Figure-SI 1.**
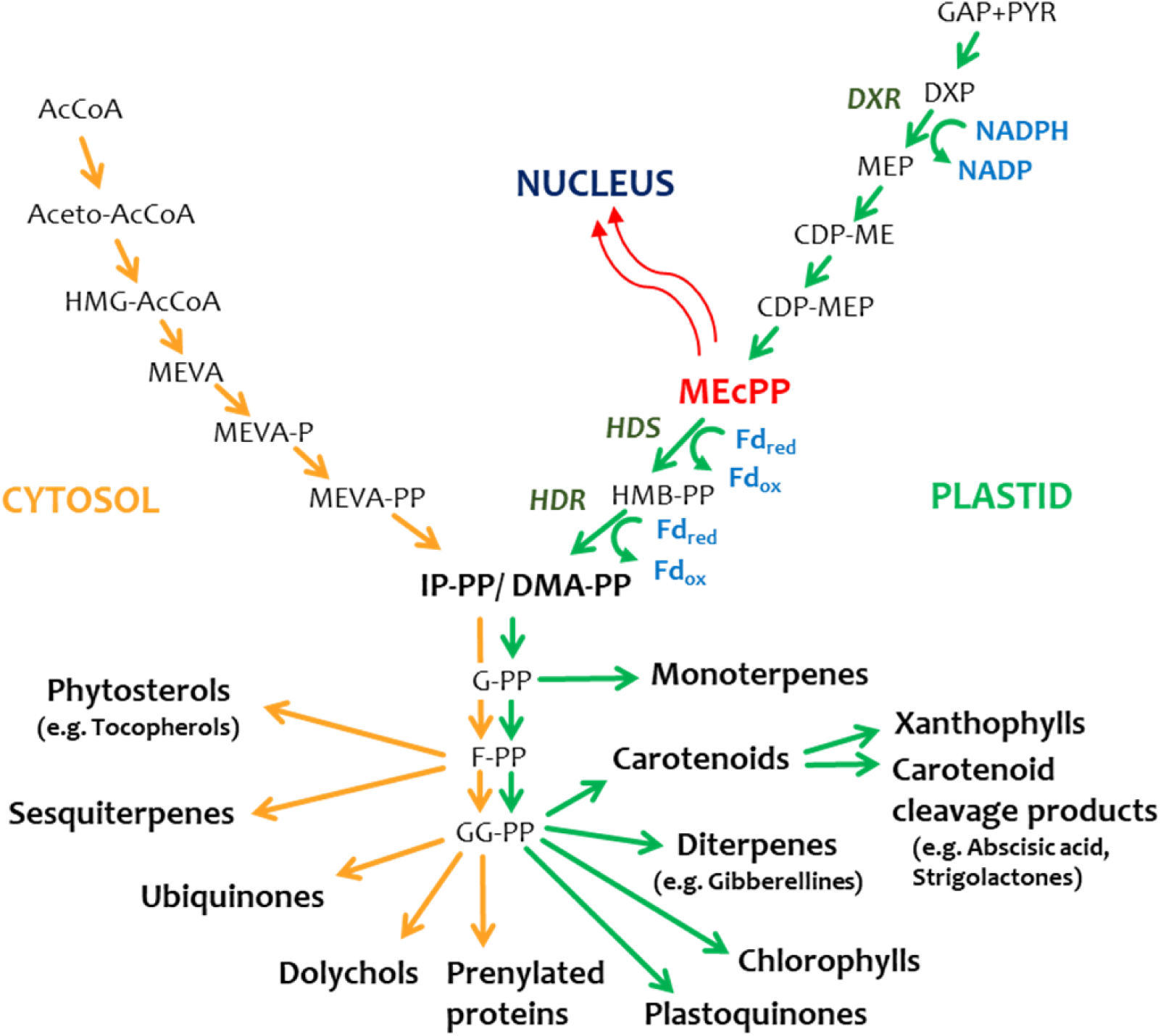
Isoprenoid biosynthesis in planta and retrograde stress signaling function of MEcPP.

**Figure-SI 2.**
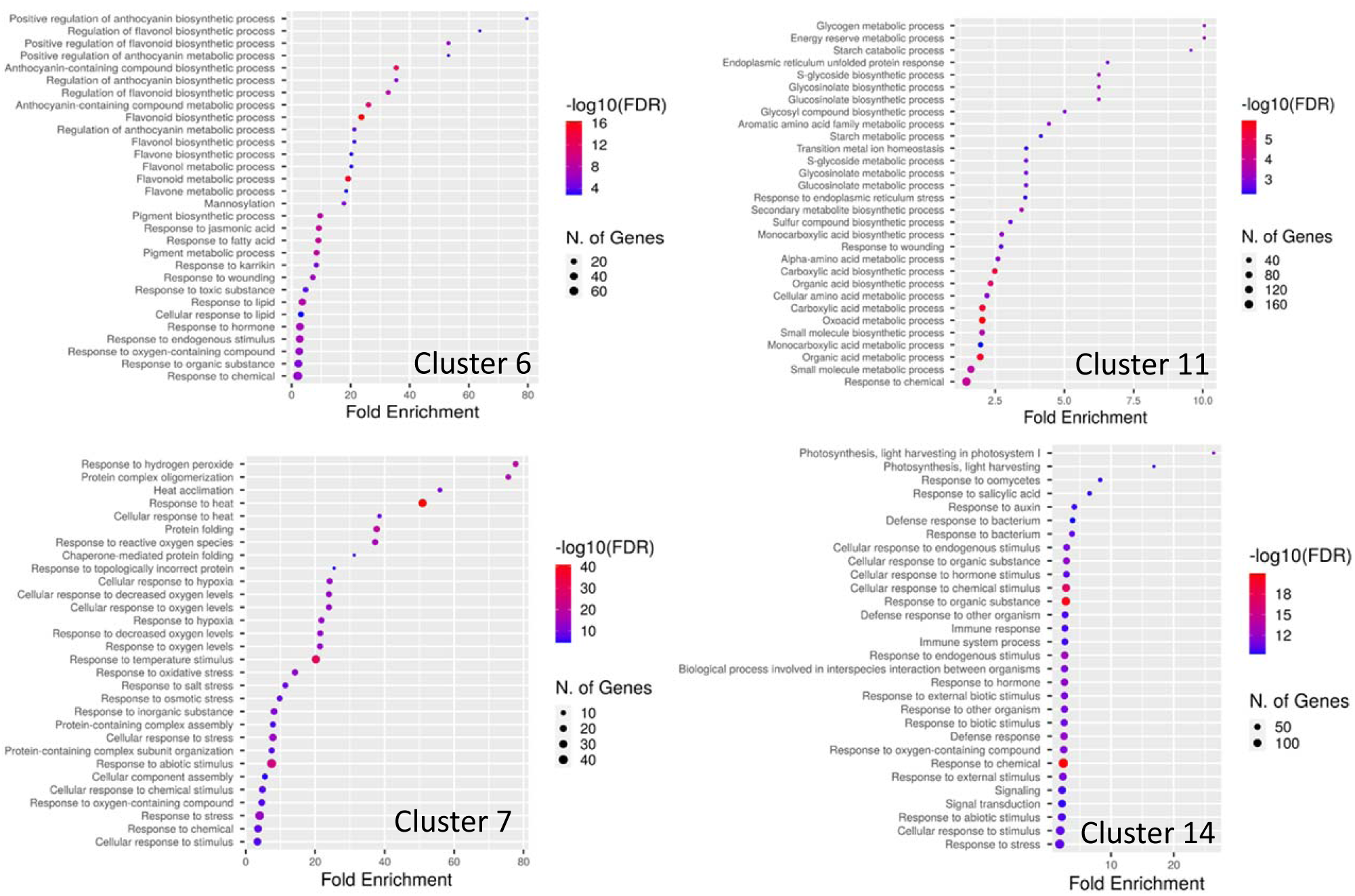
GO enrichment analysis of four selected time clusters. Time cluster analysis showing all data, all corresponding gene accession numbers, GO classes and metabolite information is listed in Table-SI 3.

**Figure-SI 3.**
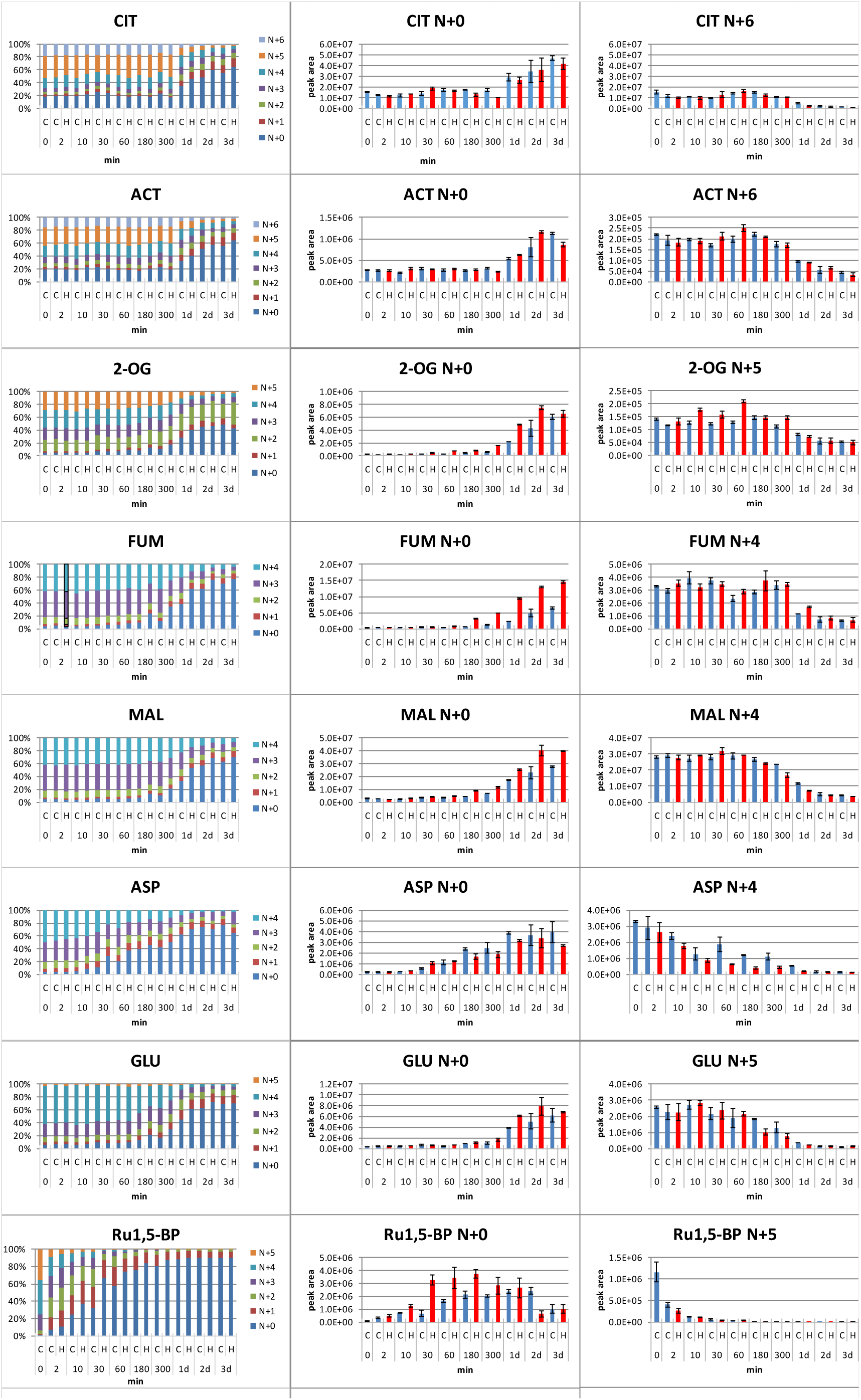
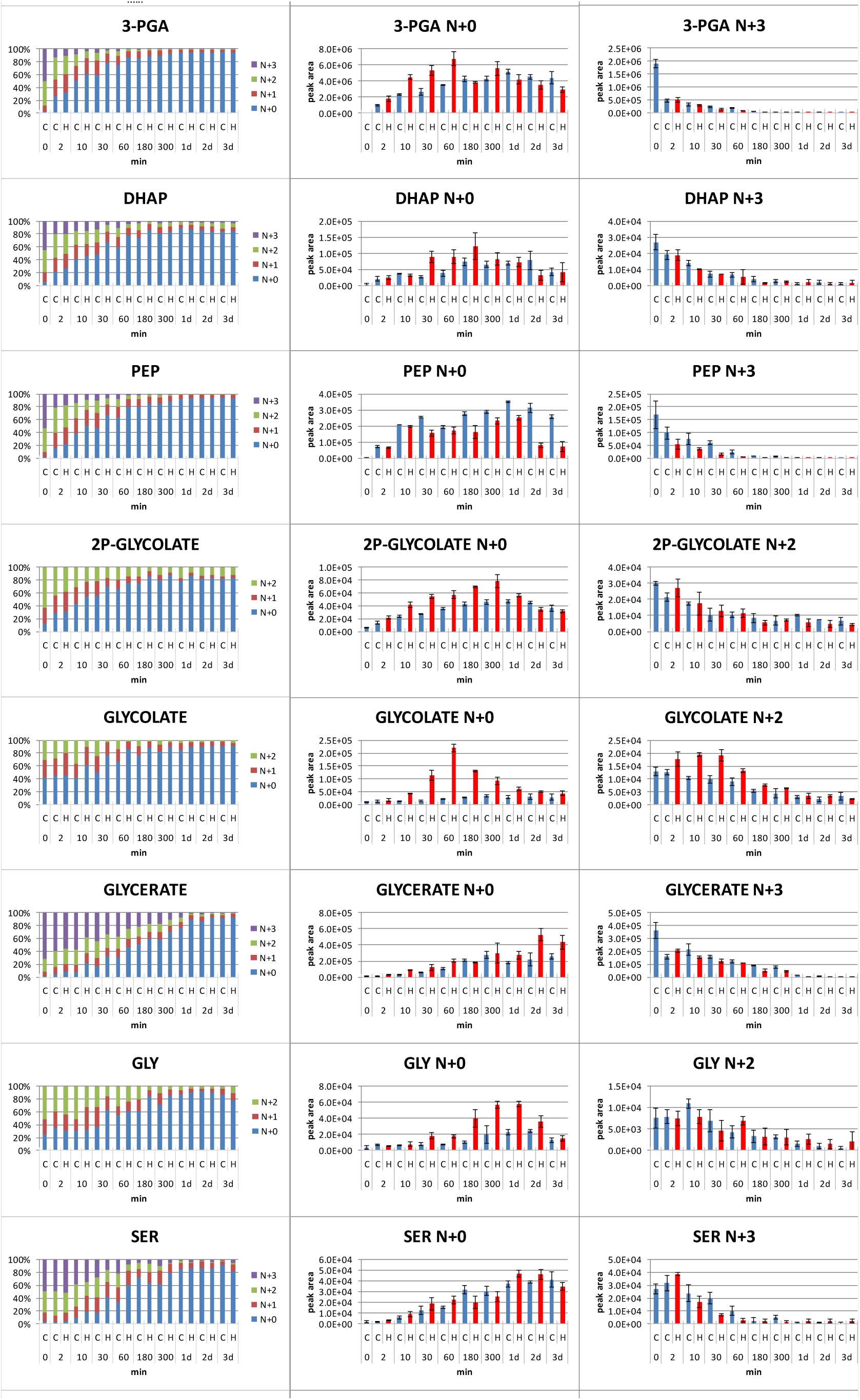
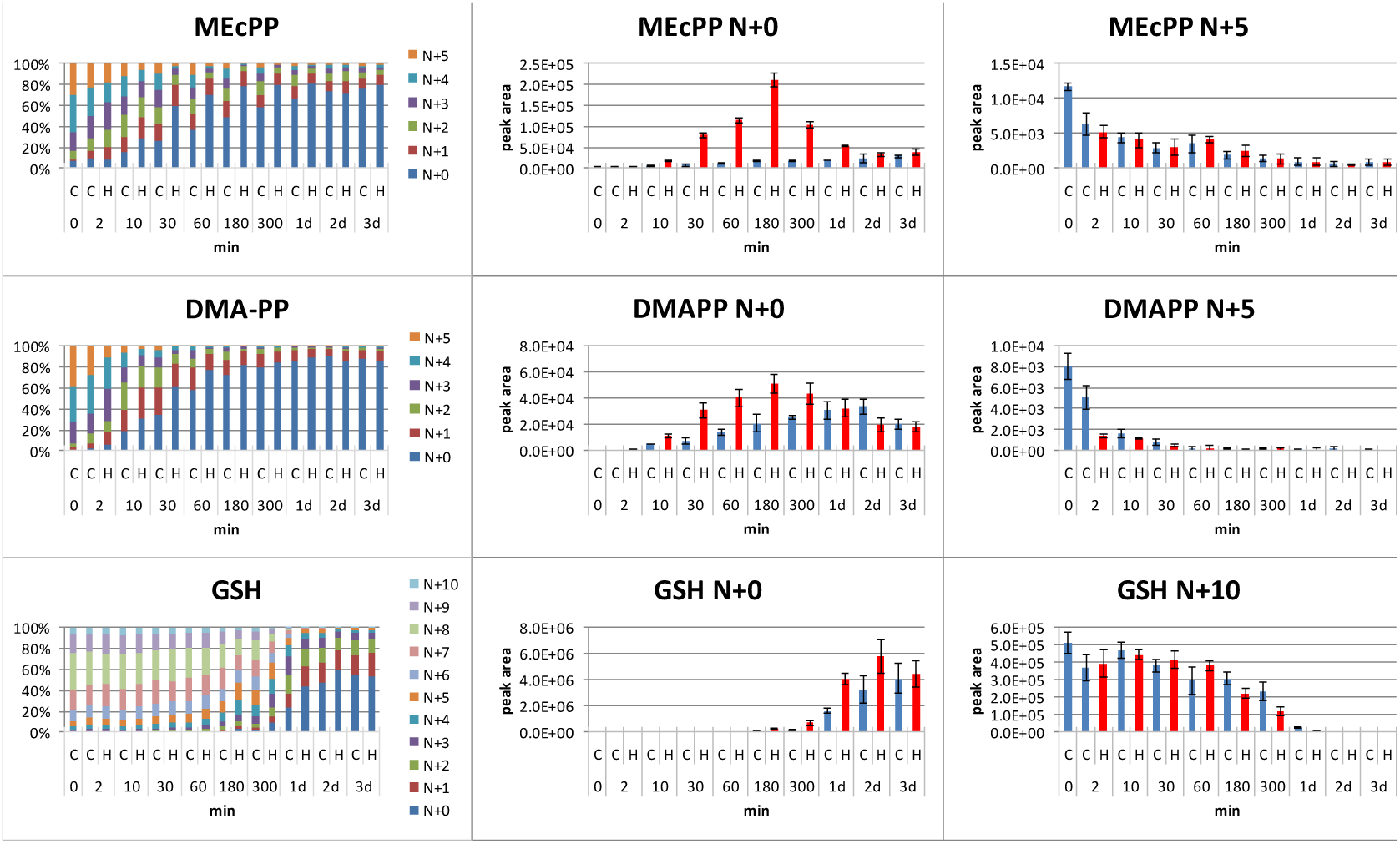
Percentage label distribution in selected metabolites (left panels) and the corresponding absolute values (as peak area) (middle panels) of the N+0 and fully labeled isotopologs (right panels).

**Figure-SI 4.**
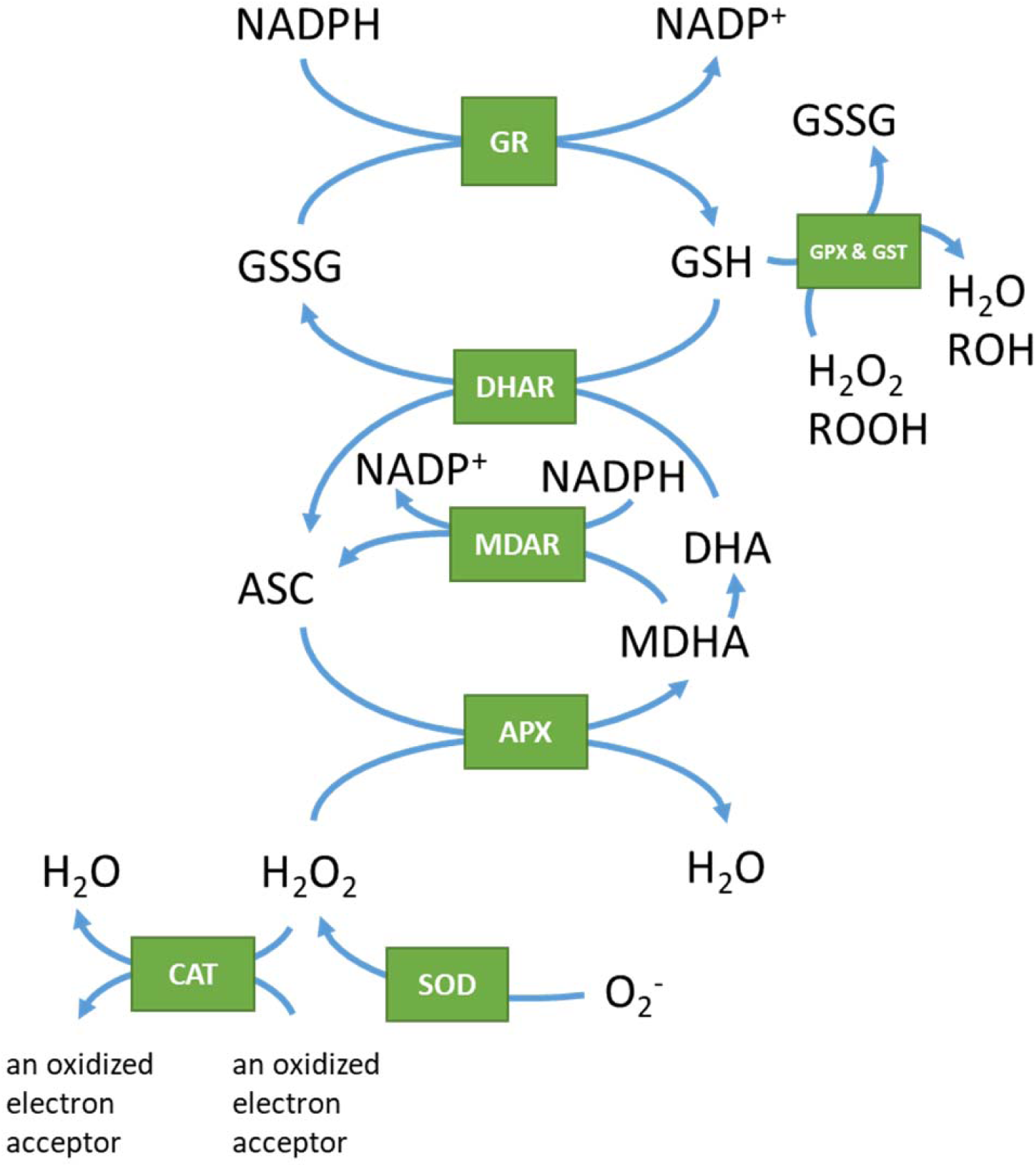
Maintenance of REDOX homeostasis: The Glutathione-Ascorbate Cycle oxidizes surplus NADPH and simultaneously reduces H2O2 and ROS. GSH oxidation is furthermore coupled to reduction of hydroperoxides.

**Figure-SI 5.**
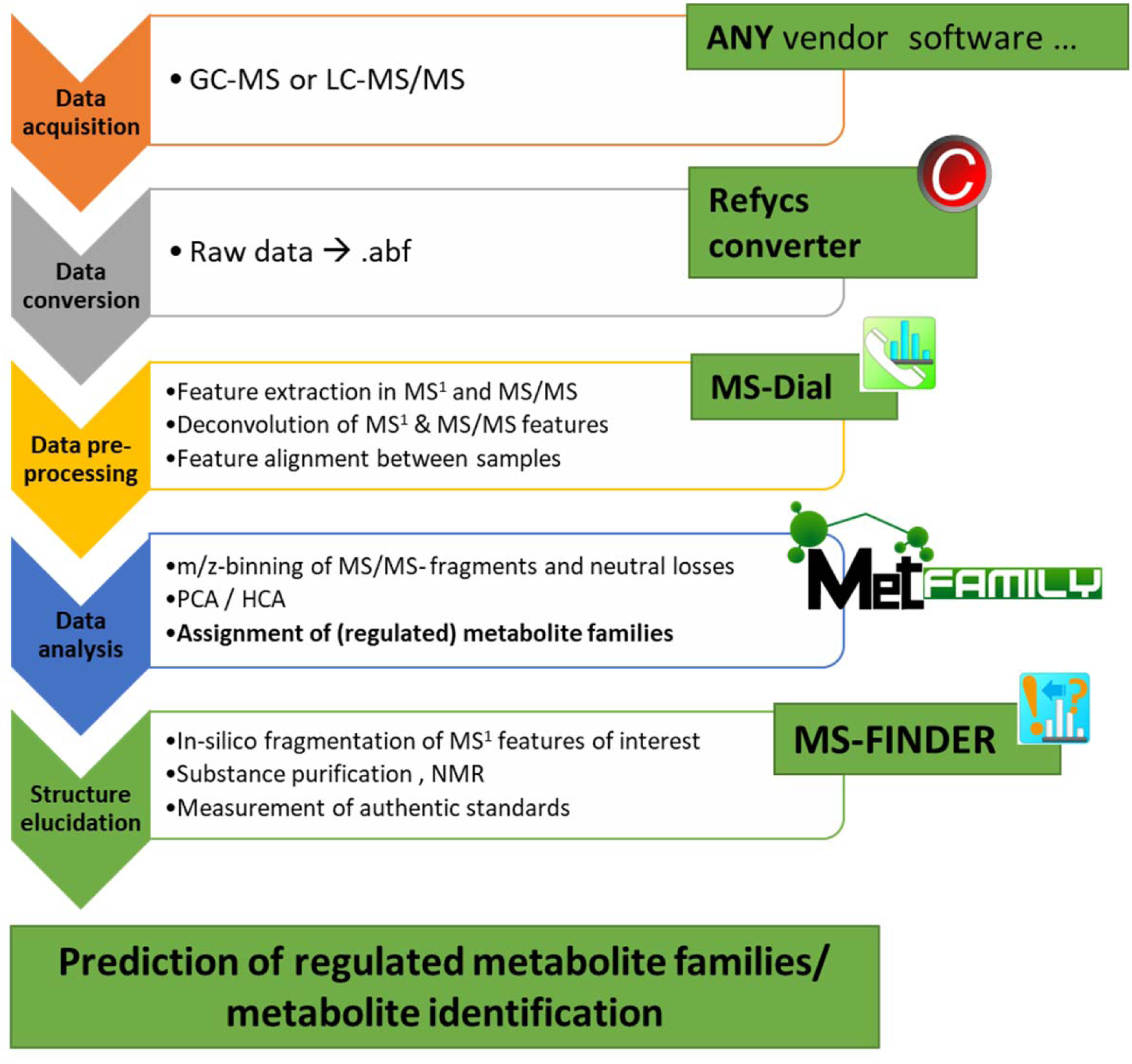
Workflow for the annotation of light-regulated metabolite families.

**Figure-SI 6.**
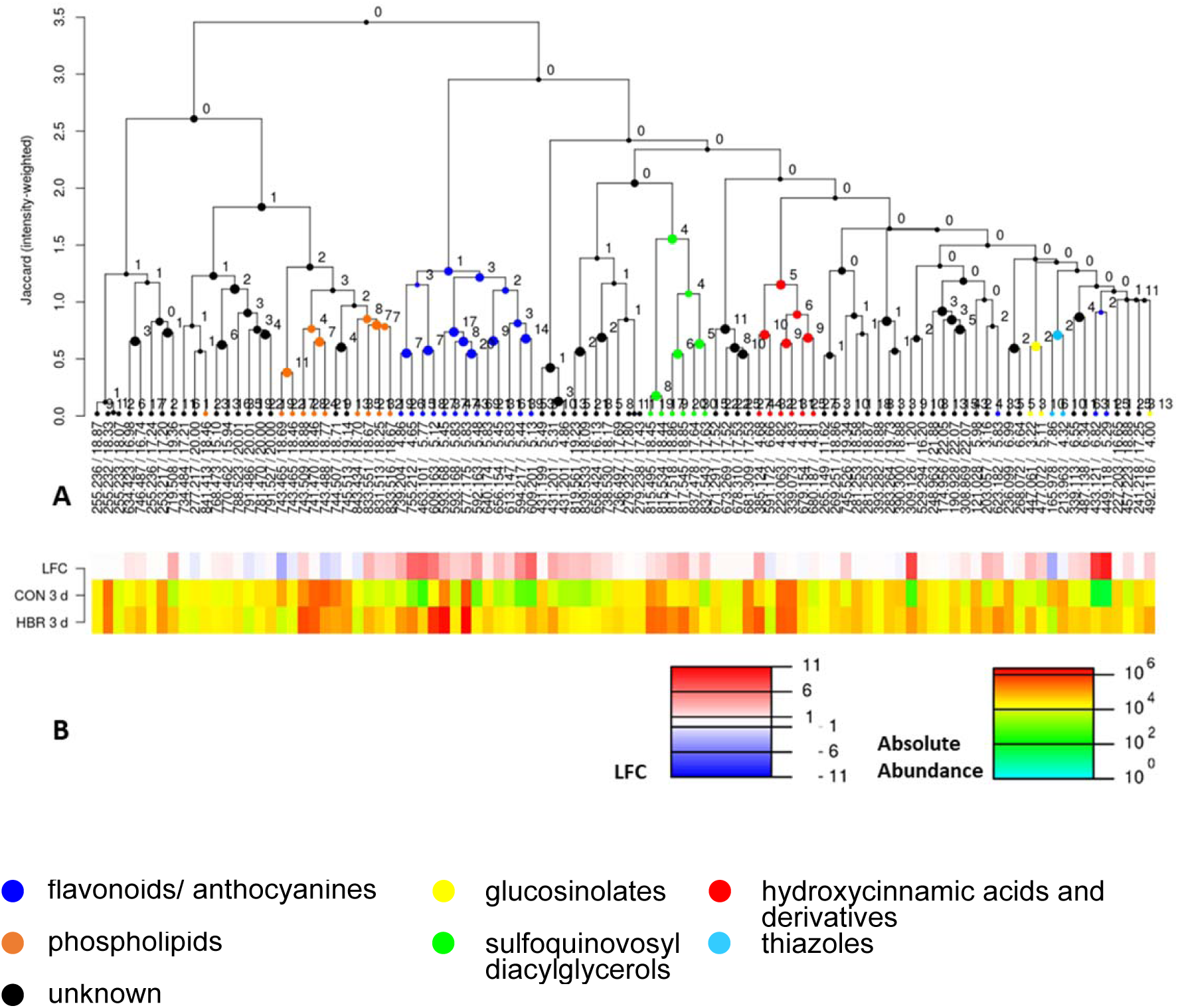
MetFamily: Hierarchical cluster analysis of semi-polar metabolites analyzed by negative mode ESI-SWATH-MS/MS. (A) MS/MS similarity dendrogram of the 90 most intense mass to retention time features, (B) average peak intensities at 72 h (n=3). Metabolite family annotations derived from molecular networking in MS-Finder, specific MS/MS features (Table-SI 7), and if available, based on best match towards MS/MS spectra of pure compounds (Lai et al, 2018; Tsugawa et al, 2019). Detail view can be obtained in MetFamily (https://msbi.ipb-halle.de/MetFamily/) by selecting the respective data sets in Data-SI 2. Structure-indicative product ions and neutral losses of specific metabolite families are listed in **Error! Reference source not found.**

**Figure-SI 7.**
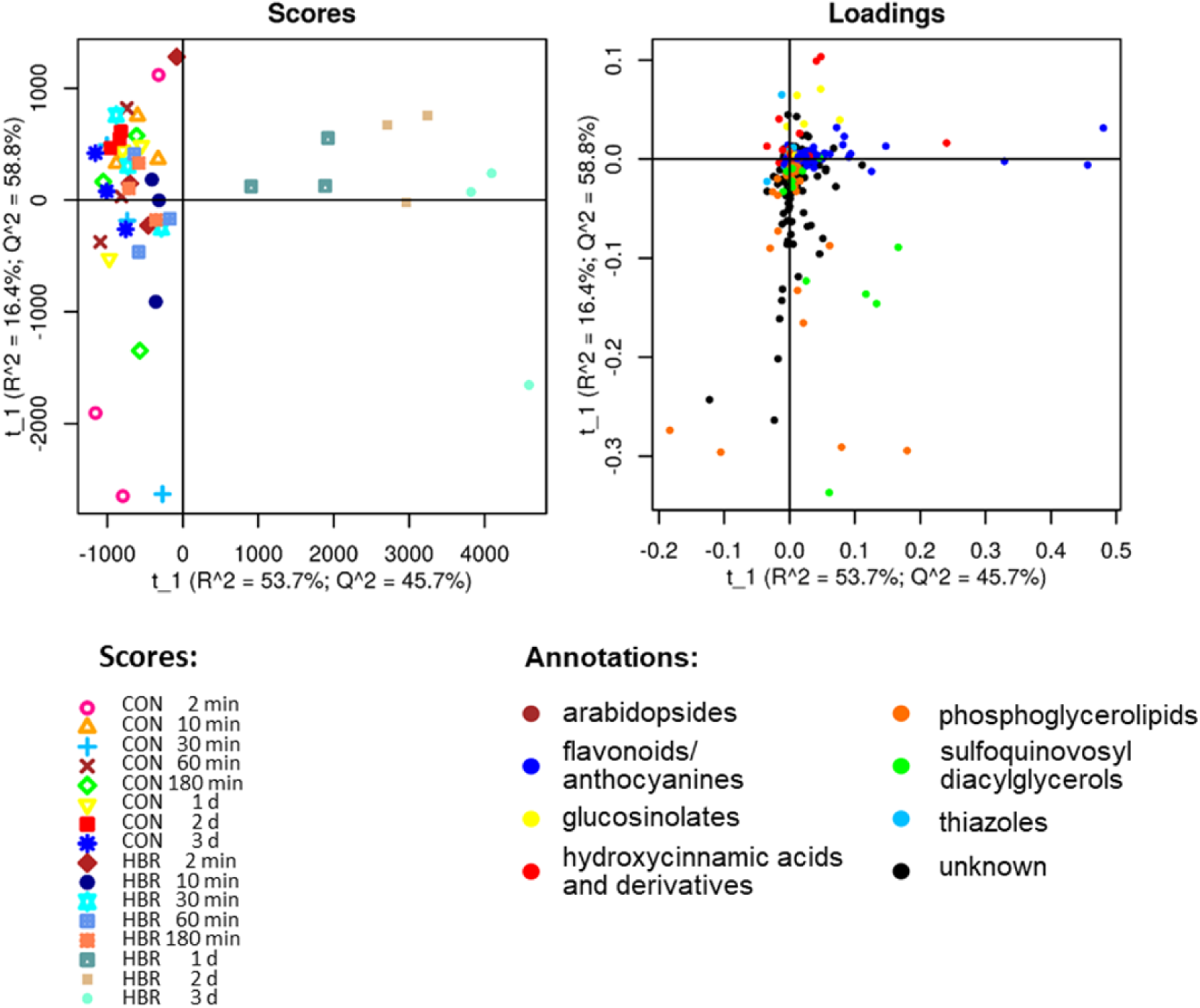
MetFamily: Principal component analysis (PCA) of semi-polar metabolites analyzed by negative mode ESI-MS, (A) Scores, (B) Loadings with metabolite family annotations (**Figure-SI 6**). Pareto-scaled PCA of 454 MS features cumulatively 59 percent of the overall variance in the data within the first 2 components. Similar analyses of separation methods 2A, 2B, 3 (see Materials and Methods) in positive mode ESI and for apolar compounds (method 4) are presented in Data-SI 2.

**Figure-SI 8.**
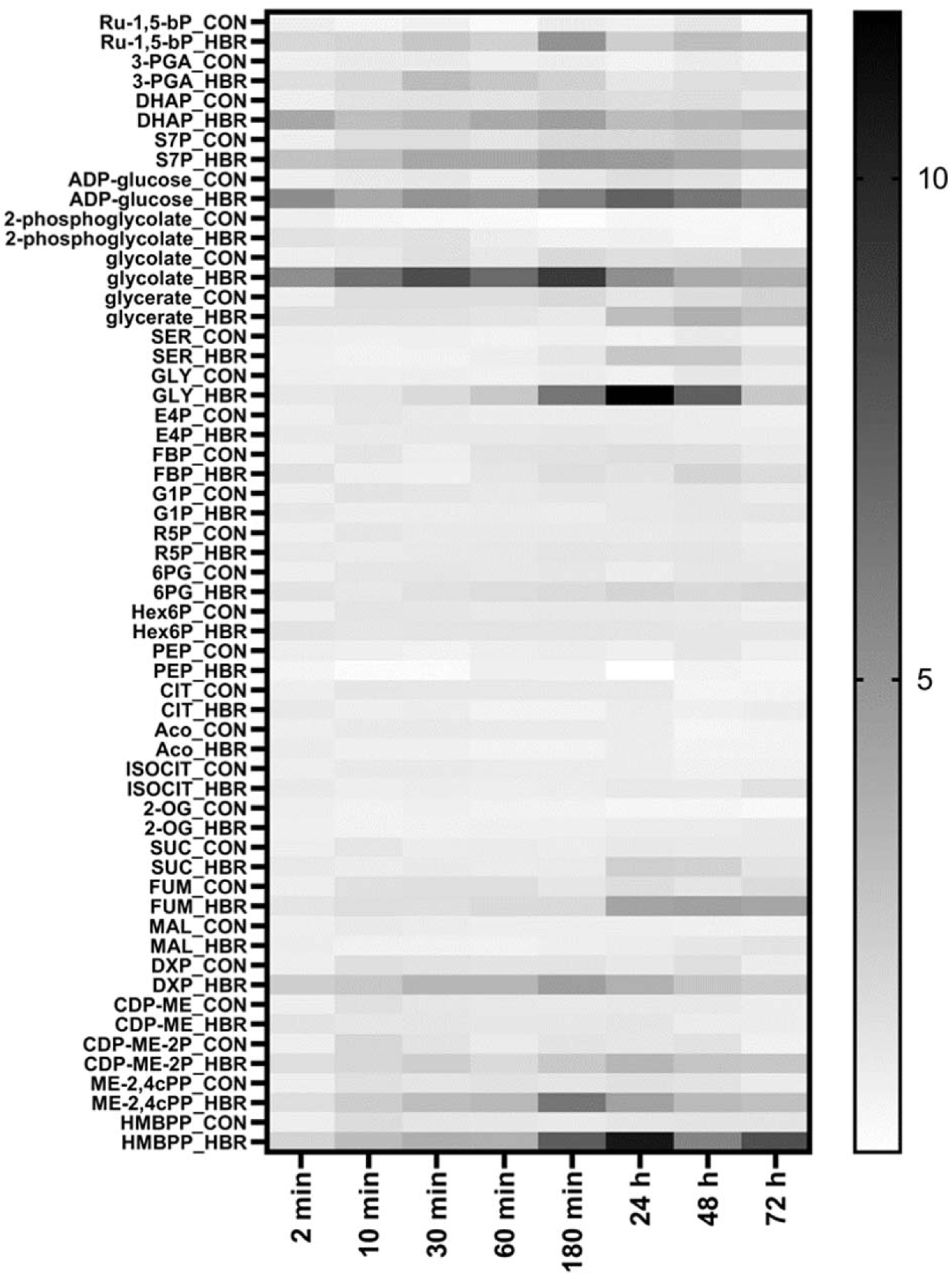
Central carbon metabolism: Normalized signal intensity of selected metabolites as shown in Figure 4 after exposure to normal light (CON) and high light (HBR); Structural identifiers for metabolite abbreviations are given in **Table-SI 5 Method parameters for the analysis of CCEM metabolites Table-SI 5**. For normalization the average peak area (n=3) of each time point was normalized to the value obtained under control light at 2 min.

**Figure-SI 9.**
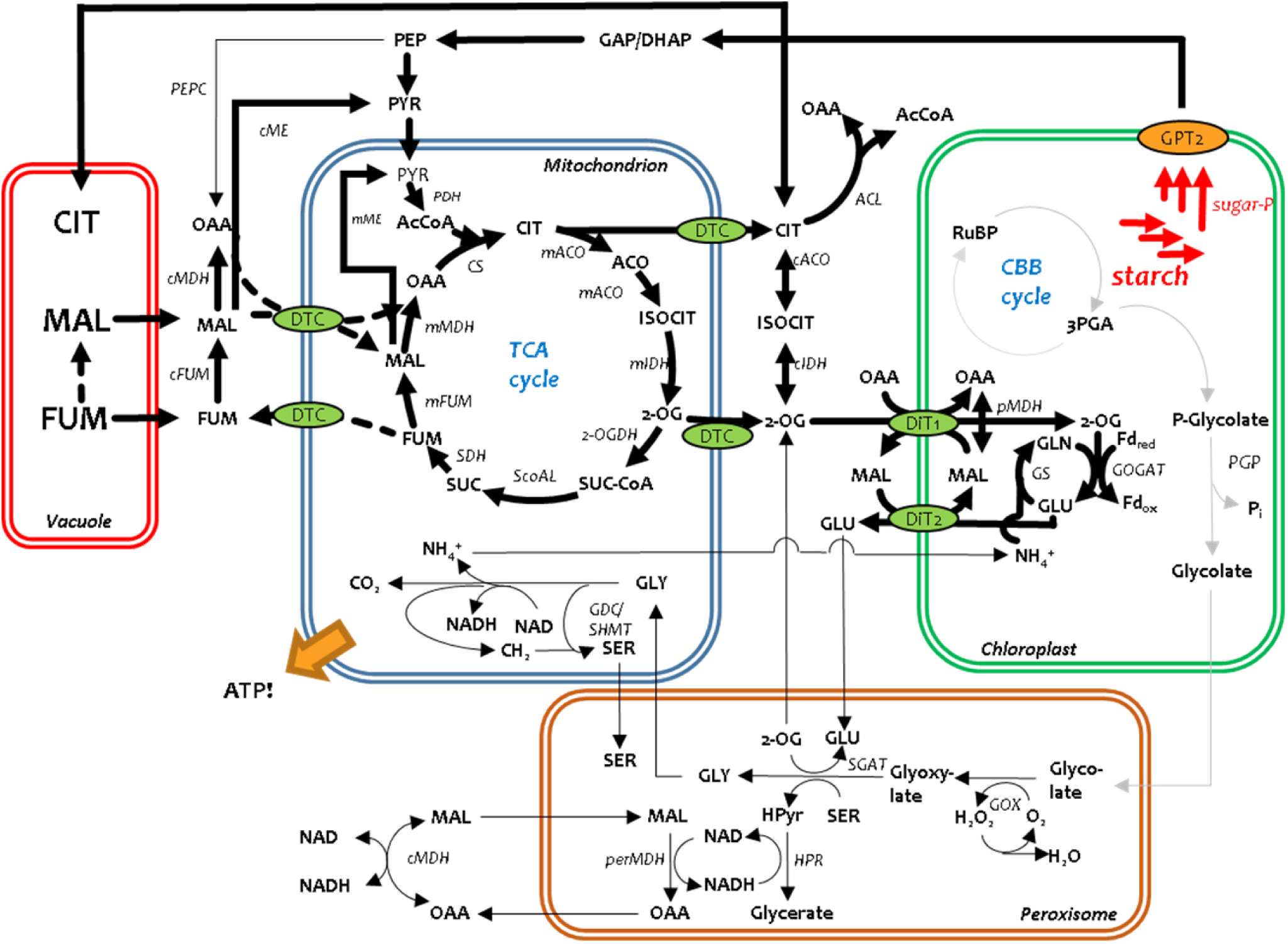
Postulated central C and N metabolism under night conditions, bold arrows symbolize fluxes, which are enhanced during the night; narrow grew arrows symbolize fluxes, which are decreased

**Table-SI 1 Redox state of SH-groups in selected peptides**

**Table-SI 2 Gene expression data of all genes.**

RNA-seq data is presented as absolute expression (tpm) and relative values, where per time point the average expression per gene of high light-stressed plants is divided by the average of the expression obtained for plants grown under control light and logarithmized to the basis 2

**Table-SI 3 K-means time cluster analysis of transcriptional regulation in response to high light stress**

**Table-SI 4 Gene expression data of individual genes of selected groups as shown in Figure 3 and Figure 8.**

RNA-seq data is presented as absolute expression (tpm) and relative values, where per time point the average expression per gene of high light-stressed plants is divided by the average of the expression obtained for plants grown under control light and logarithmized to the basis 2.

**Table-SI 5 Method parameters for the analysis of CCEM metabolites**

**Table-SI 6 Method parameters redox proteomics**

**Table-SI 7 MS/MS features used for annotation of metabolite families**

**Data-SI 1 RNA-seq NCBI Bioproject PRJNA817005 Data-SI 2** www.ebi.ac.uk/metabolights/MTBLS3109

